# Age and early life adversity shape heterogeneity of the epigenome across tissues in macaques

**DOI:** 10.1101/2025.07.13.664445

**Authors:** Baptiste Sadoughi, Rachel Petersen, Sam K. Patterson, Elizabeth Slikas, Christine Adjandba, Nicholas Ryan, Christina E. Costa, Laura E Newman, Marina M. Watowich, Cameron R. Kelsey, Ashlee Greenier, Elisabeth A Goldman, Josué E. Negrón-Del Valle, Daniel Phillips, Indya Thompson, Samuel E. Bauman Surratt, Olga González, Nicole Compo, Armando Burgos, Cayo Biobank Research Unit, Alex R. DeCasien, Kenneth L. Chiou, Christopher S. Walker, Angelina V. Ruiz Lambides, Melween I. Martínez, Kirstin N. Sterner, Lauren J. N. Brent, James P. Higham, Michael J. Montague, Michael L. Platt, Noah Snyder-Mackler, Amanda J. Lea

## Abstract

Age and early life adversity (ELA) are both key determinants of health, but whether they target similar physiological mechanisms across the body is unknown due to limited multi-tissue datasets from well-characterized cohorts. We generated DNA methylation (DNAm) profiles across 14 tissues in 237 semi-free ranging rhesus macaques, with records of naturally occurring ELA. We show that age-associated DNAm variation is predominantly tissue-dependent, yet tissue-specific epigenetic clocks reveal that the pace of epigenetic aging is relatively consistent within individuals. ELA effects on loci are adversity-dependent, but a given ELA has a coordinated impact across tissues. Finally, ELA targeted many of the same loci as age, but the direction of these effects varied, indicating that ELA does not uniformly contribute to accelerated age in the epigenome. ELA thus imprints a coordinated, tissue-spanning epigenetic signature that is both distinct from and intertwined with age-related change, advancing our understanding of how early environments sculpt the molecular foundations of aging and disease.

## Main Text

As organisms age, they experience widespread functional decline, disease, and eventually death. However, the timing, progression, and organ systems involved in these declines can vary dramatically (*1–3*). Despite growing recognition of this heterogeneity, we still lack a fundamental understanding of how aging unfolds across the body, how consistent or divergent these patterns are within and among individuals, and the degree to which environmental experiences impact variation in aging mechanisms.

A growing body of research points to social and environmental exposures—particularly adversity experienced early in life—as important determinants of age-related conditions. In both humans and other social animals, early life adversity (ELA) has been linked to age-related diseases and reduced lifespan (*4–9*). Recent evidence also suggests that ELA is associated with accelerated aging measured from blood epigenetic profiles (*10*). However, we still know little about how early life exposures shape the biological mechanisms of aging across tissues, especially at the molecular level. Addressing this gap is essential for understanding the developmental origins of age-related disease risk; however it remains difficult to investigate, especially in humans, where detailed life course information combined with multi-tissue molecular data are extremely rare.

Longitudinally monitored, long-term, and free living animal study populations are a powerful model for studies of variation in molecular mechanisms of aging and its links with ELA. While recent studies in humans and laboratory animal models have advanced our understanding of multi-tissue, age-related heterogeneity using transcriptomic and proteomic biomarkers (*1–3*, *11–18*), these studies have generally lacked naturally occurring sources of adversity and/or individual-level information about these experiences.

To address this gap, we profiled age-associated differences in DNA methylation (DNAm) across 14 tissues in 237 individuals from a well-characterized, semi-free ranging, non-human primate model: the rhesus macaques of Cayo Santiago (*19*). We focused on DNA methylation—an epigenetic modification that is one of the most studied hallmarks of aging (*20*). This mechanism is known to capture age-related molecular decay and is also responsive to environmental exposures across the life course (*21–23*). Additionally, it can be summarized into “epigenetic clocks” that provide meaningful estimates of “biological” age and that have been linked to morbidity and mortality risk (*24*). Our novel dataset integrates multi-tissue DNAm profiles with extensive data on social and environmental conditions during development (Fig. 1, A and B), allowing us to test how age and ELA impact DNAm across the body (Fig. 1C). Specifically, we hypothesized that: (i) age has tissue-dependent effects on DNAm, but that cross-tissue “biological ages” are to some degree consistent within individuals; and (ii) ELA-associated DNAm variation overlaps with age-associated DNAm differences—providing a pathway through with ELA could accelerate age-associated declines or diseases. To test these hypotheses, we first defined age-associated DNAm patterns across 14 tissues and constructed tissue-specific epigenetic clocks. We then tested the impact of six established sources of ELA (*6*) on DNAm profiles, and compared their genomic targets to those of age.

**Fig. 1.**
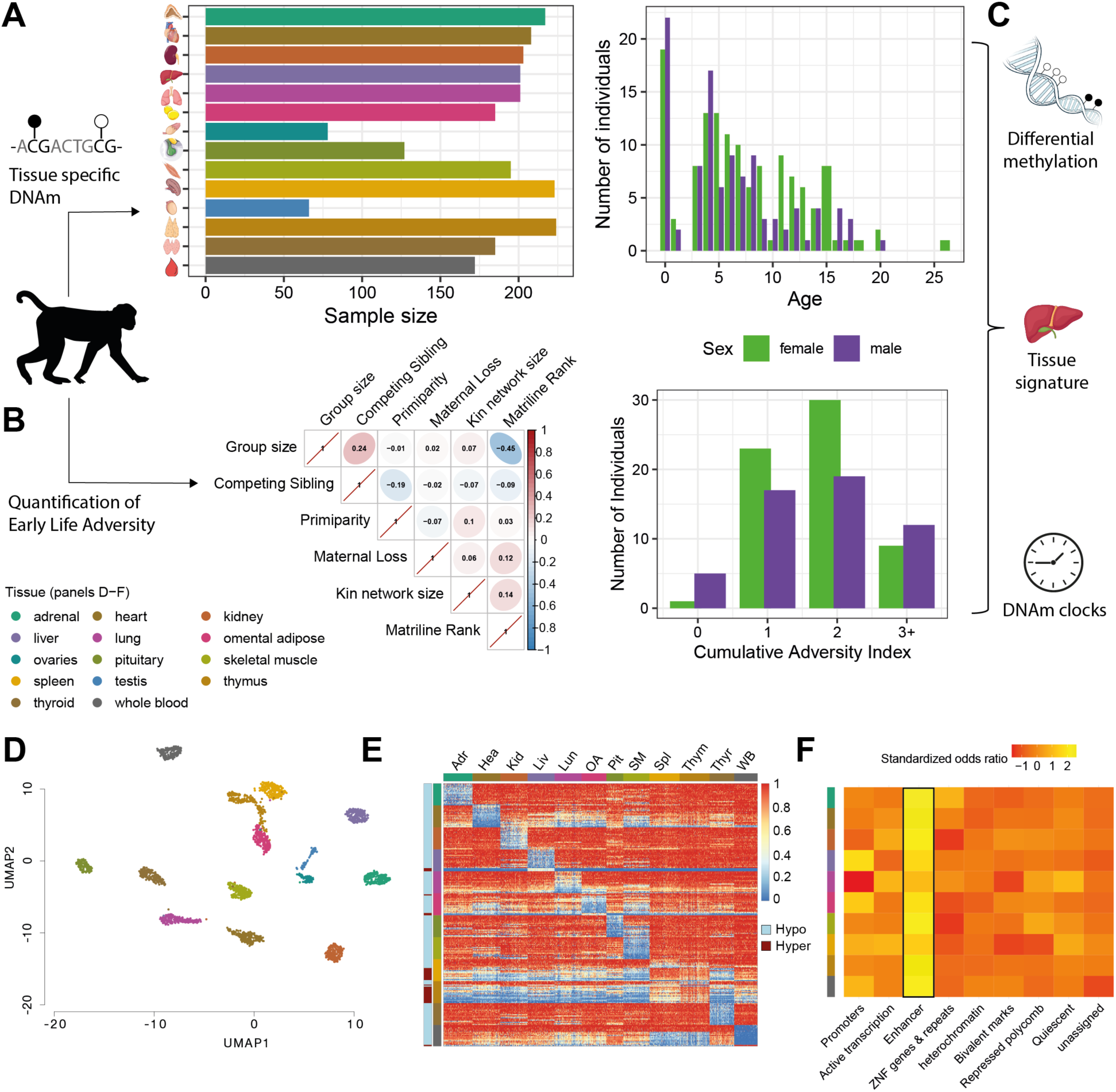
Study design on tissue-specific epigenomic variations related to age and early life adversity in rhesus macaques. (**A**) Fourteen tissues were sampled from rhesus macaques and methylation profiles generated using reduced representation bisulfite sequencing (RRBS). Tissue-specific sample size is shown with males and females plotted together for tissues other than gonads. The age range covers the entire macaque lifespan and exhibits the established female-bias in longer lifespan. (**B**) Socio-demographics traits were collected via behavioral observation and surveys. Single early life adversities (ELA) and their cumulative scores were investigated. Correlation in the occurrence of each ELA was low. (**C**) Primary response variables investigated. We tested how age and ELA impact epigenomic remodeling: (i) in specific regions of the genome, (ii) in the estimation of biological age, (iii) and on tissue-specific methylation profiles. (**D**) Dimensionality reduction of methylation profiles at 27,633 loci clusters samples according to tissue. (**E**) Top 20 tissue-specific DMRs. (**F**) Chromatin annotation of tissue-specific hypomethylated markers reveals strong enrichment for enhancers. Odds ratios were standardized by z-transformation across annotations within tissues for visualization.

### Comprehensive multi-tissue atlas of DNA methylation in rhesus macaques

We generated the largest multi-tissue DNAm dataset to date (*17*, *25*), profiling 1,309,922 CpG sites in 2,485 samples from 14 tissues sampled from 237 rhesus macaques (N = 132 females; median age = 6.12, range = 1.5 months to 25.9 years old; Fig. 1, A and C; Supplementary Materials and Methods). As expected, DNAm profiles clustered more strongly by tissue than by individual, consistent with established patterns of tissue-specific epigenetic regulation (Fig. 1D; (*26*)). We identified a total of 102,262 tissue-specific differentially methylated regions (DMRs), encompassing 681,311 CpG sites (Materials and Methods; 0.05 FDR threshold). Most tissue-specific markers (69%) were hypomethylated relative to the rest of the tissues (Fig. 1E and table S1), though the thymus stood out with almost all hypermethylated tissue-specific marks (95%). Hypomethylated, tissue-specific markers were strongly enriched near transcription start sites (TSS) and in enhancers (Fig. 1F and table S2), consistent with their role in regulating tissue-specific gene expression (*27*).

In contrast, hypermethylated markers showed less consistent enrichment for known chromatin states across tissues (table S3). Again, the thymus stood out with an overrepresentation of promoters among its hypermethylated markers (fig. S1). The thymus plays a central role in immunity and undergoes a natural process of involution with age, associated with extensive structure remodeling (*28*, *29*)—this involution likely contributes to the differences observed between the thymus’ and other tissues’ markers (see also fig. S2 and Supplemental Results). Overall, the characteristics of the tissue-specific markers we identified parallel findings in human datasets (*17*, *30*), reinforcing the evolutionary conservation of tissue-specific methylation. Moreover, this dataset represents a foundational resource for comparative epigenomics, enabling cross-tissue and cross-species analyses of DNA methylation in a well-established nonhuman primate model with translational relevance to human biology and medicine (*31*).

### Age variations in the methylome are largely tissue-dependent

To characterize the heterogeneity of age-associated epigenetic differences within and across individuals, we mapped age-associated variation in DNAm across tissues in all subadults and adult animals (≥3 years old; N = 191) using binomial mixed-effects models (*32*, *33*), controlling for sex, batch effects, social group membership, and genetic relatedness. We focused on the 12 tissues with similar sample sizes, excluding gonads (Methods). We identified 58,019 regions, comprising 652,874 CpGs, exhibiting age-associated differences in DNAm (LFSR < 0.05) (table S4). In general, increased age was associated with more intermediate DNAm levels: average DNAm was negatively correlated with age-associated effect sizes (range Pearson’s rho = −0.51 to −0.08, all p-values < 0.001, table S5). This pattern was consistent across all tissues (fig. S3), and in agreement with previous research showing that a lack of epigenetic fidelity with age tends to push both hyper- and hypo-methylated sites towards intermediate levels (*34–37*). Across most tissues, age-associated hypomethylated regions (age-hypoDMRs), where sites were less methylated with increasing age, outnumbered age-associated hypermethylated regions (age-hyperDMRs) (proportion of age-hypoDMRs to all age-DMRs in tissues: range = 0.25 - 0.94, median = 0.53). However, there was some tissue-specific variability: for example, thyroid and lung showed a strong bias towards age-associated hypermethylation (Fig. 2A and Supplemental Results). Age-associated sites (DMSs) were also enriched in chromatin states linked to gene regulation, including bivalent TSSs, enhancers, and repressed polycomb regions (Fig. 2B and table S6). Most tissues showed enrichment near actively transcribed genes, with the exception of blood (Fig. 2B). As expected (*38*), CpG islands and promoters were enriched among age-hyperDMSs, and depleted among age-hypoDMSs (fig. S4 and table S7).

**Fig. 2.**
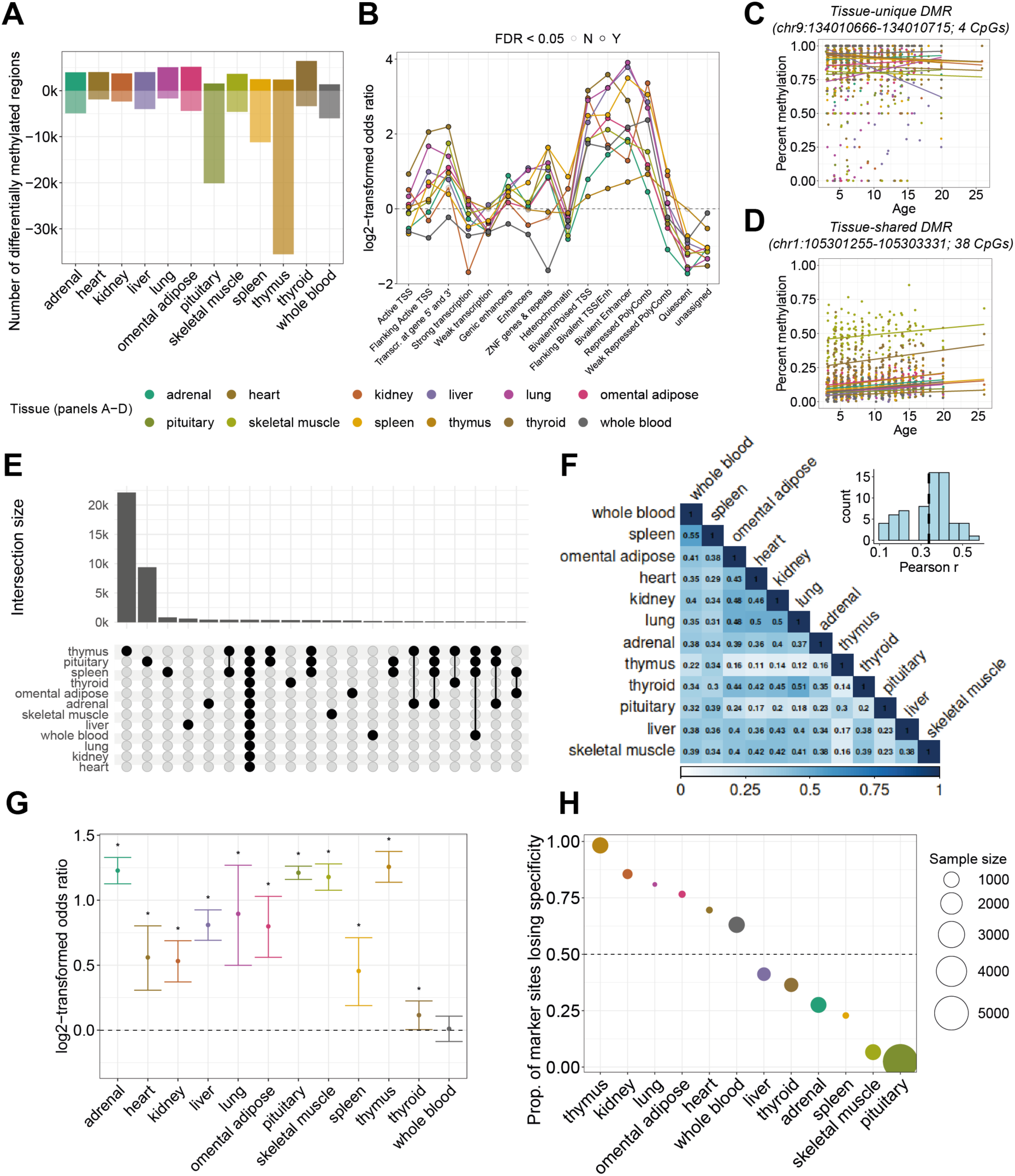
Age-associated differences in methylation levels across tissues. (**A**) Regions exhibiting a gain in methylation with age are represented with positive values on the y-axis, and regions losing methylation with age with negative values. (**B**) Enrichment for heterochromatin marks at age-associated sites in 10 tissues for which annotations were available in humans. (**C**) Percent methylation with age in one tissue-unique age-DMR, and (**D**) one tissue-shared age-DMR. (**E**) The intersection of regions exhibiting age-associated changes at LFSR < 0.05 across tissues. (**F**) Heatmap of pairwise correlations of age-associated effect sizes shows low to moderate consistency across tissues. In the histogram, the average Person’s correlation coefficient is shown by the dotted line. (**G**) Enrichment for tissue-specific markers among age-associated regions. *FDR < 0.05. (**H**) Proportion of age-DMRs that erode tissue-specificity at tissue-specific markers.

The number of age-associated regions varied substantially across tissues. The thymus and pituitary had the highest proportions of age-associated sites (>13% of tested regions), whereas the kidney and heart had the fewest (≤4%). The thymus and pituitary further stood out for exhibiting both strong and distinct age-related differences. Specifically, while “tissue-unique” age-associated regions—defined as regions significantly associated with age in only one tissue and lacking similar associations in any other tissue (Fig. 2C)—accounted for 42% of all regions tested (24,427 regions, spanning 214,154 CpGs), the prevalence of these tissue-unique regions varied widely: just 0.4% of regions in heart and blood were tissue-unique, while 29.9% and 44.2% of regions in the pituitary and thymus were tissue-unique, respectively.

On average, each individual region or CpG site was significantly age-associated in (mean ± SD) 2.5 ± 2.4 tissues (median = 1), with shared effects detected in 2.9 ± 2.6 tissues (median = 2) (Fig. 2, D and E, and fig. S5). Excluding the thymus and pituitary—the tissues that showed the greatest number of age-associated loci—did not change this result (fig. S6). Although age-DMRs shared across all tissues represented a minority of all age-associated sites (1.3%), they were significantly overrepresented relative to chance (p-value < 0.001 based on 10,000 permutations; Fig. 2E), indicating coordinated epigenetic aging in a subset of the genome. Correlations in age effect estimates between tissues were modest and variable, ranging from r = 0.11 (thymus and heart) to r = 0.55 (spleen and whole blood), highlighting the tissue-dependent nature of DNAm aging (Fig. 2F) (*39*, *40*). These results also highlight that while blood—a widely used, minimally invasive tissue—can serve as a reasonable proxy for age-associated patterns in select tissues such as the spleen, it fails to fully capture the diversity of tissue-specific patterns. Reliance on blood-based measures may thus limit insights into the complexity of molecular aging across the body (Fig. 2A).

We next tested if age-associated DNAm patterns could underlie the previously observed loss of tissue identity with increasing age (e.g., (*41*)). Tissue-specific epigenetic markers (Fig. 1E) were significantly enriched for age-associated DMRs (Fig. 2G and table S8), suggesting that aging can alter the very regions that define tissue-specific methylation patterns. On average, 12.4% ± 10.9% of tissue-specific markers were also age-associated DMRs, ranging from 4.7% in blood to 42.7% in the thymus. However, these age effects did not uniformly erode tissue identity. In half of the tissues, tissue-specific methylation became less distinct with age, while in others, such as the pituitary, it became more pronounced (Fig. 2H and fig. S7). For example, kidney-specific hypomethylated markers tended to gain methylation in older individuals—a pattern also observed in aging human kidneys and linked to declining renal function (*42*). In the thymus, the apparent erosion of tissue identity likely reflects widespread epigenetic remodeling associated with age-related thymic involution (Fig. 2, A and B). In contrast, the pituitary exhibited greater epigenetic distinctiveness at increased ages. Together, these findings show that age can either blur or sharpen tissue-specific methylation profiles, depending on the tissue. This nuanced pattern mirrors results from recent multi-tissue gene expression studies in humans and mice (*41*, *43*), highlighting the likely variable impact of aging on molecular tissue identity.

### Tissue-specific epigenetic clocks capture heterogeneity in age

To quantify within and between individual heterogeneity in molecular age, we developed tissue-specific epigenetic clocks (Fig. 3A, fig. S8, and table S9 (*24*); Materials and Methods). These clocks were highly accurate: we could predict the chronological age of a held-out sample to within 1.1 years within a given tissue (median absolute error range: 0.82 in omental adipose to 1.53 in ovary; Fig. 3B and table S10), and predicted ages were strongly correlated with chronological age (ranging from r = 0.81 in the ovaries to 0.95 in the liver; Fig. 3B, fig. S9, and table S11). We confirmed our clocks’ reliability by applying the whole blood clock to additional Cayo rhesus macaque samples not included in this study (Pearson’s rho = 0.76, MAE=2.11; Supplementary Material and table S11). From the tissue-specific clocks, we regressed individuals’ clock-predicted ages on their chronological age and extracted the residuals—these values are referred to as age deviations (*44*) and reflect individual-level measures of acceleration (or deceleration) of biological aging within that tissue (table S12 and fig. S10).

**Fig. 3.**
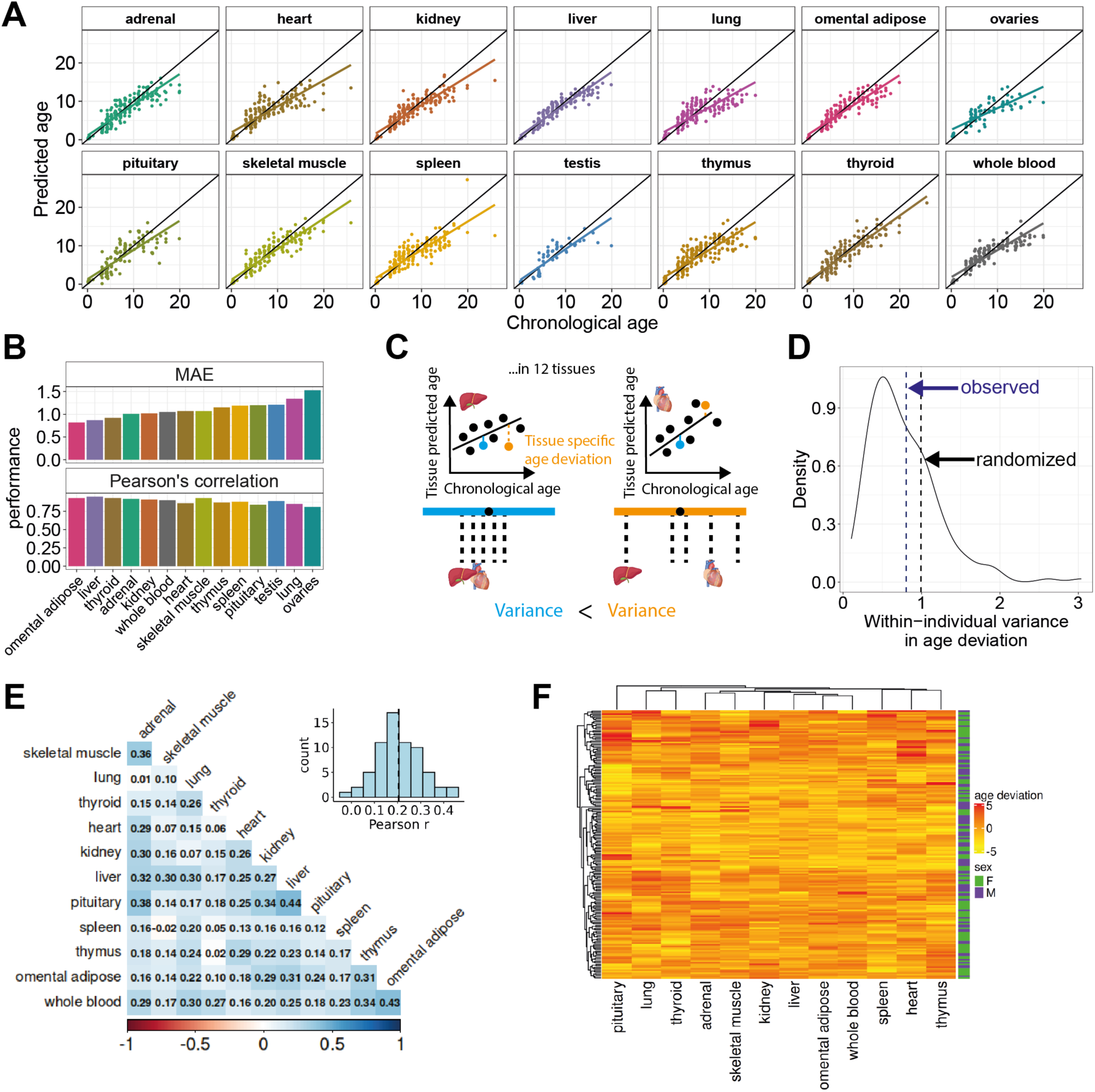
Organ specific age predictions and within-individual heterogeneity in methylation age. (**A**) Scatter plots of predicted ages based on methylation profiles in 14 organs. The diagonal depicts x=y and the colored regression lines the best linear fit. (**B**) ‘Clock’ model prediction accuracy is given by the Median Average Error (in years) and the Pearson correlation coefficient with chronological age. Organs are ordered on the x-axis by increasing MAE. (**C**) An example of within-individual variance in DNAm ages across organs from two individuals: one exhibiting low variance (blue, i.e., low within-individual heterogeneity) and one exhibiting greater variance (orange, i.e., high within-individual heterogeneity). (**D**) The intensity of within-individual heterogeneity varies between individuals (density plot), although on average (dotted blue line) the level of within-individual heterogeneity is lower than could be expected by chance (the average of 10,000 randomizations calculating the mean across individuals is depicted by the dotted black line) consistent with a degree of within-individual consistency. (**E**) Pairwise correlation coefficients between age deviations across organs. (**F**) Within-individual and between-individual heterogeneity in age deviations across 12 organs. Age deviations exhibit low correlations across organs which lead to an absence of consistent within-individual, multi-tissue profiles (i.e., ageotypes).

Using these tissue-specific age-deviations, we asked if individuals showed similarity in their age-deviations across all tissues, or if there was instead no evidence for within-individual consistency in biological aging. To do so, we used two complementary statistical approaches. First, we found that the average within-individual variance in age deviations was significantly lower than a randomized null model in which these analyses were repeated but tissue samples were shuffled across individual identity (p < 0.001; Fig. 3, C and D). Second, we used linear mixed effects models to estimate that individual identity explained 19% of the variance in age predictions (controlling for chronological age, sex, group, and tissue type; LRT = 182.11, p-value < 0.001; table S13). Both approaches support the observation that age deviations were weakly to moderately positively correlated across most tissue pairs (Fig. 3E), such that individuals tend to look ‘old for their age’ or ‘young for their age’ across most tissues in their body. However, we found no evidence for distinct subgroups of individuals—so-called “ageotypes”—who consistently aged faster or slower across the same distinct subset of tissues (Fig. 3F, fig. S11, and Supplemental Results). In other words, patterns of aging heterogeneity are generalized and not easily clustered into consistent, coordinated tissue sets at the epigenetic level, a conclusion on par with the most recent evidence from proteomics (*14*).

Next, we tested if demographic factors might influence the degree of within-individual consistency we observed. Males and females showed similar levels of within-individual heterogeneity (effect size ± SD = −0.03 ± 0.07, p = 0.7). However, within-individual heterogeneity of predicted age increased with chronological age (effect size ± SD = 0.106 ± 0.033, p = 0.002; fig. S12). This relationship was not statistically significant when restricted to older individuals (>6 years). This could reflect a biological constraint, namely that the degree of inter-tissue epigenetic divergence may be established early in life, with individual differences in aging trajectories becoming entrenched and relatively stable over time. To investigate how tissue-specific patterns may be shaped early in life, we next drew on comprehensive demographic and behavioral records to examine the impact of early-life adversity on DNAm patterns and epigenetic age across tissues.

### Early life adversities shape DNA methylation at distinct loci with cross-tissue consistency

We compiled six established, ecologically relevant sources of ELA (N = 191 per ELA, unless stated otherwise): maternal loss, maternal primiparity, limited maternal kin network, low matrilineal rank (N = 117), presence of a close-in-age younger sibling (N = 190), and being born into a large social group. For individuals with complete data (N = 116; Materials and Methods), we also derived a cumulative adversity score by summing exposures across the six individual sources (median: 2, range: 0-4). These individual ELA sources, and cumulative ELA metric, reflect social or environmental conditions previously linked to survival in this and other non-human primate populations (*6*, *8*), and parallel known adversity exposures in humans (*45*, *46*).

Because these ELA sources were largely uncorrelated (Fig. 1B), unlike sources of adversity in humans that often co-occur (*47*), we could assess the independent and joint effects of distinct adversities. Using binomial mixed-effects models, we separately tested for the effect of each ELA, as well as cumulative ELA, on DNAm. In an effort to isolate the effects of early-life environment, we included age, sex, social group, and adult dominance status (an influential aspect of the current social environment) as covariates. Across tissues, we identified 7,533 DMRs (containing 198,740 CpG sites) significantly associated with at least one ELA (LFSR < 0.1); this ranged from 2,235 to 95,469 ELA-associated CpG sites per tissue (Fig. 4A and table S14). Additionally, we identified 785 DMRs (containing 14,058 CpG sites) significantly associated with cumulative ELA, ranging from 5-6,114 CpG sites per tissue. Following the same criteria as we used for understanding effect size sharing in age-associated sites (Fig. 2E), we found evidence for broad sharing of a given ELA’s effects on DNAm across tissues—both in terms of which sites were targeted (Fig. 4B and fig. S13) and the direction of the ELA effect (Fig. 4C and fig. S14). In contrast, each individual ELA was associated with DNAm at a variable number of CpG sites (Fig. 4D), and within a tissue, each ELA generally targeted distinct CpG sites (Fig. 4E and fig. S15). Together, these results suggest that different ELAs may impact distinct molecular pathways and processes, but that the epigenetic response to any given ELA is to some degree coordinated across the body.

**Fig. 4.**
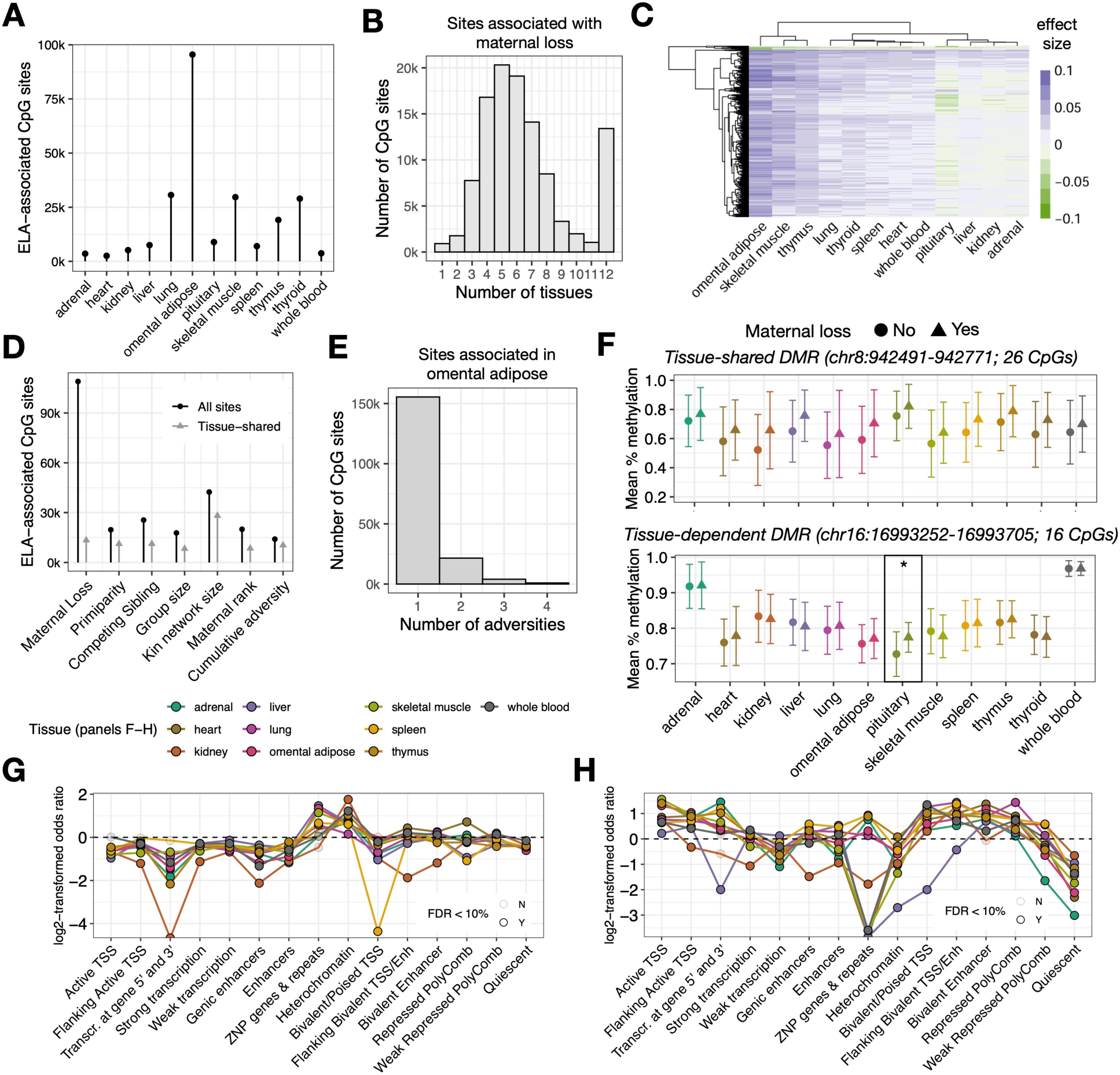
Early life adversity effects on DNA methylation across tissues. (**A**) Number of CpG sites per tissue that were associated with at least one form of ELA. (**B**) The intersection of CpG sites associated with maternal loss across tissues following the effect size sharing analysis. (**C**) Hierarchical clustering of sites associated with maternal loss, colored by effect size (positive = increased methylation, negative = decreased methylation), limited to sites that did not have shared effects across all 12 tissues. (**D**) Number of CpG sites associated with each ELA in at least one tissue. (**E**) The intersection of CpG sites associated with each ELA in omental adipose tissue. (**F**) Mean percent methylation per tissue for a representative tissue-shared DMR (top), and tissue-dependent DMR (bottom). Circular points represent mean percent methylation for individuals who did not experience maternal loss, triangle points represent individuals who did experience maternal loss, asterisk denotes which tissue exhibited the ELA effect in the tissue-dependent example. (**G**) Enrichment of tissue-shared and (**H**) tissue-dependent ELA-associated CpG sites within different chromatin states.

### Early life adversity targets regulatory regions and generates tissue-specific vulnerability

DMRs exhibiting shared responses to ELA across all tissues were enriched in canonically less active chromatin states, such as heterochromatin (OR = 1.1–3.4, FDR < 0.001; Fig. 4, F and G, and table S15). In contrast, sites with tissue-dependent effects (i.e., sites that were ELA-associated in some but not all tissues) were enriched in more active regulatory regions, such as active transcription start sites (OR = 1.2–2.9, FDR < 0.001; Fig. 4, F to H, and table S15). These results suggest that while many ELA effects are consistent across all tissues, tissue-dependent ELA effects may be most prone to impact gene regulation.

We next tested if certain tissue types are more susceptible to ELA. Grouping tissues by cell lifespan and biological function (table S16, Supplementary Methods), we found that tissues with long-lived cells exhibited stronger ELA effects than those with faster cell turnover (linear model: estimate = –0.014, SE = 0.003, p = 2.6 × 10⁻7; fig. S16 and table S17). Immune and endocrine tissues were also affected more than metabolic and other tissues (immune vs. metabolic: estimate = 0.014, SE = 0.003, p = 2.1 × 10⁻5; endocrine vs. metabolic: estimate = 0.018, SE = 0.002, p < 2 × 10⁻¹⁶; fig. S17 and table S17), highlighting the biological systems most likely to mediate long-term effects of adversity. These findings align with previous research showing ELA effects immune on function and hormonal regulation (*48*).

### Early life adversity mirrors the effects of age in a tissue-dependent manner

Finally, we asked whether ELA alters the same pathways or regulatory elements as those affected by age. If so, ELA could accelerate or otherwise reprogram key features of the aging methylome, thus implicating ELA-altered DNAm as a conduit through which early life experiences shape age-related morbidity and mortality patterns. To test this possibility, we examined the extent to which ELA recapitulates age-associated methylation patterns across tissues.

Focusing on epigenetic age deviations from our tissue-specific epigenetic clocks, we found that experiencing ELA was not significantly associated with age acceleration across tissues (Fig. 5A and table S18). Within tissues, the only significant association between ELA and age acceleration was observed in the pituitary following maternal loss (estimate = 1.49, SE = 0.54, FDR-adjusted p-value = 0.05; fig. S18 and table S19). Yet, ELA-associated sites were strongly enriched among age-associated sites across tissues and adversities (Fig. 5B and table S20), suggesting that ELA and aging impact similar genomic targets. However, the only tissue where age and ELA showed similar directional effects (e.g., where ELA exposure was similar to older age) was the pituitary (OR = 3.3, FDR = 1.19 × 10⁻5, Fig. 5C and table S21), whereas immune tissues (spleen and thymus) showed significantly discordant effects of age and ELA (thymus: OR = 0.43, FDR = 0.004; spleen: OR = 0.59, FDR = 0.08; Fig. 5C and table S21).

**Fig. 5.**
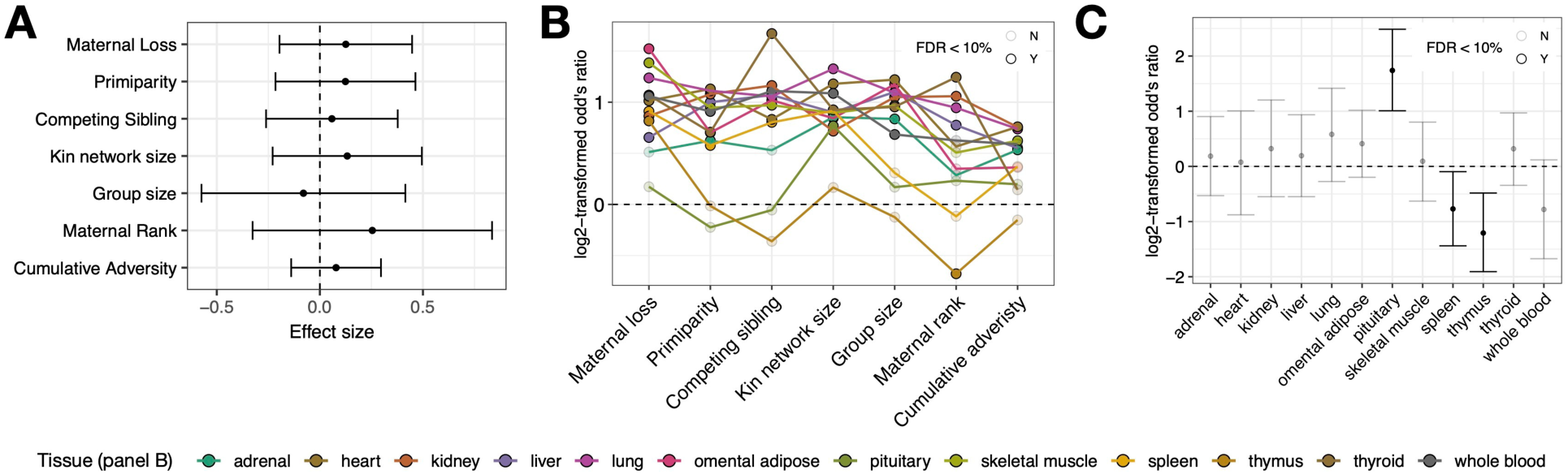
Early life adversity contributes to age-related DNAm variation. (**A**) ELA was not significantly associated with age acceleration across tissues. Points represent model estimates and error bars represent the standard error. (**B**) Enrichment of ELA-associated sites among age-related sites across tissues, and (**C**) enrichment for consistent direction of age and maternal loss-associated differences in DNAm (Fisher’s exact test).

## Discussion

We generated an extensive multi-tissue DNA methylation dataset in a semi-free ranging, well-characterized nonhuman primate population, and revealed how age and ELA intersect to shape the epigenome across the body. This dataset not only fills a critical gap in comparative primate epigenomics, where large multi-tissue datasets are absent outside of humans, but it also allowed us to address fundamental biological questions about tissue-specific patterns of aging and their environmental influence.

The effect of age on DNA methylation varied markedly across tissues, with approximately half of the tissues exhibiting a loss of identity—a presumed cause for functional decline with age (*41–43*). Despite heterogeneous age effects across tissues, tissue-specific epigenetic age estimates remained coordinated within individuals—pointing toward environmental factors that accelerate or decelerate biological aging across the organism. These findings echo insights from plasma proteomes (*14*) and provide a reason why tissue-specific diseases can be predicted, albeit with lesser accuracy, from clocks derived from other tissues (*1–3*, *16*, *49*). In contrast to the tissue-dependence of age effects, the epigenetic signature of early adversity was surprisingly synchronized across tissues. However, different types of adversity targeted largely non-overlapping sets of CpG sites, suggesting advantage in one domain (e.g., high maternal social status) may not be able to offset or buffer the biological consequences of adversity in another (e.g., maternal loss). Overall, the contrast between the tissue-dependent effects of age and ELA suggest that while aging involves more localized and tissue-contingent remodeling, early adversity may act through more systemic mechanisms.

This work points to several future directions. It will be important to address why certain tissues exhibit age-related loss of tissue identity while others do not, and to understand the connection between this methylation erosion and tissue-specific functional capacity and disease. As methylomes were generated from bulk samples, the relative contributions of tissue-specific changes in cell composition as opposed to intra-cellular remodelling of the methylation landscape remains to be determined. Further, determining which adversities have the most enduring effects, and which might instead decay over time, will be important for understanding under which circumstances ELA may contribute to accelerating aging. Longitudinal studies are especially important for ruling out the possibility that selective disappearance might influence our and other cross-sectional results. To address these questions in the absence of confounds and limitations faced by human studies (*10*), the establishment of deeply behaviorally and biologically phenotyped naturalistic animal models hold particular promise (*50*, *51*). Our results underscore this need to integrate developmental context to uncover the roots of heterogeneity not only in disease risks, but in the fundamental pace and patterning of biological aging.

## Acknowledgments

We thank the Caribbean Primate Research Center for their invaluable contribution to the long-term data collection and maintenance of the Cayo Santiago population. We thank Olga Konstantara and Ryan Rossow for their assistance with tissue preparation and DNA extractions. We thank all members of the Lea and Snyder-Mackler labs for their thoughtful feedback on this work.

## Author contributions

Conceptualization: NSM, AJL, MJM, JPH, LJNB

Methodology: BS, RP, NSM, AJL, SKP, ES, CA, CEC, LEN, MMW, KLC, CRK, AG, EAG, KNS, MJM

Validation: BS, RP, NSM, AJL, MMW, ES, CA, AG

Formal analysis: BS, RP Investigation: BS, RP, SNL, AJL

Resources: NSM, AJL, JPH, LJNB, MLP, AVRL, MJM

Data collection: JENDV, DP, IT, SEBS, OG, NC, AB, CBRU, ARD, CSW

Data curation: BS, RP, NSM, AJL Visualization: BS, RP, NSM, AJL

Funding acquisition: NSM, AJL, MJM, LJNB, JPH, MLP, AVRL, MIMM

Project administration: NSM, AJL, MJM, LJNB, JPH, MLP, AVRL, MIM

Supervision: NSM, AJL

Writing – original draft: BS, RP, NSM, AJL Writing – review & editing: all authors

## Data and materials availability

Genomic sequences generated as part of this study have been deposited in NCBI’s Sequence Read Archive under accession number PRJNA1286127. Bioinformatic and statistical scripts are available from github https://github.com/BaptisteSadoughi/CayoCrosstissueDNAm.

## Supplementary Materials

Cayo Biobank Research Unit Scientific Stakeholders

## Materials and Methods

Supplemental Results

Figs. S1 to S18

References

Other Supplementary Materials for this manuscript include tables S1 to S21 in excel format.

