## Supplemental Methods and Figures for "Age and early life adversity shape heterogeneity of the epigenome across tissues in macaques"

**The PDF file includes:**

Cayo Biobank Research Unit Scientific Stakeholders
Materials and Methods
Supplemental Results
Figs. S1 to S18
References

**Other Supplementary Materials for this manuscript include the following:**

Tables S1 to S21 in Excel format

### Materials and Methods

Study design

We studied the behavior and collected the postmortem tissues of rhesus macaques living on Cayo Santiago, Puerto Rico (18°09 N, 65°44 W). The macaques freely-range on the island and are organized into social groups which include several adult females and males mirroring the species natural social organization. Female rhesus macaques are usually considered adults from the age at first birth or around 5 years old. Males undergo a first growth spurt and are fully mature around 8 years of age but achieve reproductive maturity earlier. Behavioral observation and census surveys were used to determine individuals' exposure to ELA. As part of population control measures (*52*), two entire social groups were removed in 2016 and 2018, and necropsies were conducted by the CPRC veterinary staff. All samples come from this database. All procedures related to the removal of animals were conducted by the CPRC following standard protocols approved by the Institutional Animal Care and Use Committee (IACUC) at the University of Puerto Rico (Protocol #338300). The sample size was 132 females and 105 males (Fig. 1, C and D).

Tissue collection and preservation

Sedation was performed by the administration of ketamine (100 mg/kg body mass). Immediately after, blood was drawn via femoral venipuncture and 4 mL was collected using BD Vacutainer® K2-EDTA collection tubes. Following veterinary euthanasia, complete diagnostic necropsies were performed, whereby several tissues were extracted and weighed for complete documentation of disease state and histopathology. Tissues investigated for this study included the adrenal (cut on transverse plane including medulla and cortex), heart (near the perfusion site on the right ventricle), kidney (right kidney, after removal of the capsule), liver (on the right lobe), lung (on the middle lobe of the right lung), omental adipose tissue (on the first incision of the peritoneal cavity), ovary (cut on coronal plane), pituitary (collected whole), skeletal muscle (collected from the quadriceps at anterior belly of rectus femoris or vastus lateralis from the right leg), spleen (from the superior angle of the body nearest to the tail of the pancreas), testes (small piece of lobule after removal of the tunica), thymus (collected closer to trachea to limit collection of heart adipose tissue), and thyroid (whole left thyroid). Small biopsies of tissues were collected and placed into pre-labeled vials (with necropsy ID printed on a cryolabel along with sample ID and tissue type) containing 0.5 ml of RNAlater, ensuring the tissue was completely submerged. Once all tissues were collected from an animal, the tubes were placed in sample boxes and stored at 4°C overnight to ensure the RNAlater had saturated the tissue. After 24 hours, the storage boxes were moved to a -20°C freezer for no more than 1 week and then stored at -80°C until extraction.

DNA extraction, bisulfite conversion, sequencing, and data pre-processing

We extracted DNA using the Zymo Quick-DNA/RNA MagBead kit (Zymo Catalog No. R2131), following the manufacturer's instructions for animal tissue and dual DNA/RNA extraction. Prior to extraction, approximately 0.1g of tissue was excised and homogenized by bead beating, using a Qiagen Tissuelyzer II. After homogenization, we followed the manufacturer's protocol for dual DNA and RNA extraction. For a subset of the blood samples, extraction from whole blood was performed at the University of Washington, Seattle in 2019 using the QIAamp Fast DNA Tissue Kit (QIAGEN), following the manufacturer's instructions.

DNam was quantified by reduced representation bisulfite sequencing (RRBS) using 300 ng of DNA as input (*53*). RRBS bisulfite conversion was performed using the EZ-96 DNA Methylation-Lightning MagPrep kit (Zymo Research), and then libraries were ligated with unique dual index sequencing primers and amplified with 16 cycles of PCR. We combined RRBS libraries in equimolar quantities and sequenced them on an Illumina NovaSeq. Given that the fragment sizes generated by RRBS are often shorter than the read length, we only used read 1 from each read pair to avoid double-counting of the same nucleotide.

<u>Data pre-processing, quality check and libraries selection</u>
Single-end FASTQ files were trimmed using *Trim Galore!* (function trim_galore) https://www.bioinformatics.babraham.ac.uk/projects/trim_galore/ and aligned in *bismark* (*54*) (functions bismark, --score_min L,0,-0.6 -R 10 \ -p 4) to the rhesus macaque reference genome (Mmul_10). Cytosine methylated and total counts were extracted with functions bismark_methylation_extractor and coverage2cytosine (flag --merge_CpG). We assembled count data using the package *bsseq* (function read.bismark) (*55*) for further processing and analysis in RStudio version 4.4.0 (*56*).

Based on quality checks (QC), we selected libraries with a number of uniquely mapped reads between 2.5M and 100M, a ratio of uniquely mapped reads to total reads >0.35, and a conversion rate < 0.02 at CHH loci. Libraries matching best to their respective whole genome were retained (18 libraries from infants best matching their mothers were also included). CpG sites with coverage in < 0.25 of the samples and median coverage < 5 were removed. CpGs within 1000 base pairs of each other were clustered into regions, and regions constitutively hypomethylated (average percent methylation < 0.1) or hypermethylated (average percent methylation > 0.9) across all fourteen tissues were removed. We filtered out samples with less than 0.70 of regions covered based on visual quality check. Finally, we applied tissue-specific filtering removing regions with coverage < 0.25 in the tissue to ensure robust tissue-specific inferences.

To identify possibly mislabelled libraries, we tested our ability to retrieve tissue identity from the overall methylation profile using two unsupervised clustering techniques and a multinomial classifier. We used Uniform Manifold Approximation and Projection implemented in the package *umap* (*57*) with n_neighbors=50 and min_dist=0.5 and other parameters to default, and Principal Component Analysis with prcomp(). Samples were colored according to their tissue of origin. We considered libraries as mislabelled when they failed to cluster with their respective tissue on the UMAP, the PCA, and when the correct tissue identity could not be predicted with more than 0.5 certainty by a multinomial elastic net regression in *glmnet* (*58*). Excluded libraries were replaced by duplicates when available, and the entire QC procedure was conducted again until no samples were mislabelled. The final sample size was 2485 across 14 tissues (Table 1).

**Table 1.** Tissue-specific cohort details.

| Tissue | Mean age | Median age | Min age | Max age | N° female | N° male |
|---|---|---|---|---|---|---|
| adrenal | 6.83 | 6.13 | 0.12 | 19.93 | 117 | 100 |
| heart | 6.85 | 6.13 | 0.12 | 25.88 | 108 | 100 |
| kidney | 7.12 | 6.20 | 0.12 | 25.88 | 113 | 90 |
| liver | 6.63 | 6.08 | 0.13 | 19.89 | 110 | 91 |
| lung | 6.56 | 6.08 | 0.12 | 19.93 | 102 | 99 |
| omental adipose | 6.75 | 6.08 | 0.12 | 19.93 | 102 | 83 |
| ovaries | 7.53 | 6.39 | 0.20 | 19.91 | 78 | 0 |
| pituitary | 6.95 | 6.20 | 0.12 | 19.89 | 70 | 57 |
| skeletal muscle | 7.09 | 6.15 | 0.12 | 25.88 | 106 | 89 |
| spleen | 6.88 | 6.12 | 0.12 | 25.88 | 125 | 98 |
| testis | 5.80 | 5.18 | 0.24 | 19.93 | 0 | 66 |
| thymus | 6.81 | 6.06 | 0.12 | 19.93 | 122 | 102 |
| thyroid | 6.86 | 6.08 | 0.13 | 25.88 | 103 | 82 |
| whole blood | 6.92 | 6.14 | 0.12 | 19.91 | 96 | 76 |

Relatedness

We generated whole genome sequencing data from DNA extracted tissue samples using either the Nextera DNA library preparation kit and paired-end sequencing on the Illumina HiSeq 2500 or using the Illumina DNA preparation kit and paired-end sequencing on the Illumina NovaSeq (NovaSeq6000 or NovaSeqX). Raw reads were mapped to the macaque rheMac10 reference genome (*59*) using the command-line function 'fq2bam' (*60*). Genotypes were imputed with 'glimpse2' (*61*) using a population-specific reference panel. Imputed genotypes were filtered for an imputation quality score > 0.9, leaving 14,447,329 SNPs in the final VCF files, which were then used to generate a genetic relatedness matrix using the program KING (v2.3).

Early life adversity

We used long-term data cultivated by the CPRC to calculate six unique sources of ELA, all of which may influence an infant's ability to receive adequate social and/or nutritional support. These ELA sources have been well established in other non-human primate models of ELA (*8*, *62*, *63*), have been associated with survival in this study population (*6*), and are homologous to sources of adversity in humans (*64*).

1. Maternal loss: The death of a mother, particularly during lactation but also after weaning, significantly increases an infant's risk of mortality (*65*). We define maternal loss as the death of a mother—whether from natural causes (disease or injury, n= 33) or permanent removal from the island (n= 6)—within the first four years of life, a critical developmental period before individuals transition from juvenility into adulthood, during which maternal loss has been shown to significantly impact offspring survival in primates (*9*). While natural death likely reflects a period of maternal decline and may impose a

greater degree of adversity, the removal of a healthy mother may still disrupt offspring support. Given the profound consequences of both, we classify these experiences together.

2. Primiparity: Offspring of primiparous mothers face a heightened risk of mortality (*66*), suggesting that first time mothers may not provide comparable social and/or nutritional support as more experienced mothers.
3. Maternal kin network: Familial support, often measured by kin network size, is important for survival in cercopithecine primates, including this species (*67*). We define maternal kin network size as the number of closely related adult females present in the group on an individual's date of birth, serving as a proxy for familial support. To ensure biologically meaningful connections, we limit our kin networks to individuals with a relatedness coefficient $\geq 0.063$ (mothers, grandmothers, sisters, and aunts)–the threshold at which individuals in this population recognize kin vocalizations (*68*). We focus specifically on maternal kin as both males and females typically interact more frequently with their maternal relatives than with other groupmates (*69*).
4. Matrilineal rank: Female rhesus macaque social hierarchies are generally structured by matrilines, wherein all individuals within a matriline rank above or below all individuals from another matriline. Dominance rank strongly influences resource access (*70*) and can impact offspring nutritional support (*71*, *72*). CPRC staff determined matriline ranks annually within each social group, categorizing them high, medium, or low based on dyadic agonistic interactions. Because matriline members share rank relative to other matrilines, this serves as a proxy of individual social status. Matriline ranks remain stable over time (*73*), allowing us to extrapolate rank from adjacent years in instances where matriline rank data were missing, following (*6*).
5. Competing sibling: Infants with siblings close in age may receive reduced maternal investment, and a short interbirth interval is linked to increased offspring mortality in this study population (*74*). We defined close-in age siblings as those born within 355 days after the focal individual's date of birth–representing the bottom quartile of IBIs in this population (*6*). We did not consider the IBI preceding the focal individual's birth and thus last-born offspring could not experience this adversity.
6. Group size: Large group sizes are associated with feeding competition and reduced reproductive rate in primates and other taxa (*75*, *76*). We used group size as a proxy for within-group competition, defined as the number of adult individuals (male or female) in the group on an individual's date of birth.

All ELA sources were recorded as a binary variable: presence=1, absence=0. For group size and kin network size, which were recorded as the number of individuals within the group/ kin network, we considered individuals born into the top quartile of group sizes (> 103 individuals) and the bottom quartile of kin network sizes (< 2 individuals) as having experienced adversity. Additionally, we considered individuals born into low ranking matrilines as having experienced adversity. We took the sum of these six measures to approximate cumulative ELA, as cumulative

ELA scores have been shown to be better predictors of fitness proxies in humans and non-human primates (*5*, *8*, *77*).

Current social status

Social status at time of sampling, included as a control variable, was assigned based on behavioral observations following methods previously described (*78*), and categorized into high (top 20% ranks), medium (20 to 80% ranks), and low (lowest 20%) sex-specific tiers. When behavioral observations were insufficient to assign rank for young individuals (n = 127 typically < 6 years of age), we relied on known maternal relatedness (*67*, *79*). Individuals in their natal groups were assigned the rank of their mother if she was still alive. If the mother was deceased, the rank of first-degree relatives (such as sisters or aunts) was assigned, provided that all female relatives were confirmed to belong to the same rank class. Additionally, three adult males who had recently migrated to their new group were classified as low rank, as males are known to queue for dominance in this species and population and thus would enter a new group as the lowest rank (*80*).

Tissue-specific markers

To identify tissue-specific markers, we focused on DNA methylation at a subset of 179,969 regions (representing 1,281,603 CpG sites) covered in all 12 tissues (excluding gonads). We conducted one-versus-all comparisons with the MACAU binomial mixed models algorithm implemented in *PQLSEQ* (*32*, *33*). The models included age, sex, a relatedness matrix, and percent uniquely mapped reads in the sample to account for batch and other technical effects. Models that failed to converge were excluded. Loci were deemed tissue-specific when they exhibited significant differential methylation of at least 20% fold-change with all other tissues in a consistent direction of change. To gain insight into the functional relevance of identified markers and permit comparisons with human atlas, we annotated tissue-specific markers with seven-state chromatin marks based on chromHMM 15-state descriptions (see also below).

Differential methylation with age and early life adversity

We tested for age-associated changes in methylation levels using binomial mixed effects models implemented in *PQLSEQ* (*32*, *33*). Due to the large number of infants, we decided to exclude very young individuals below three years of age from our analysis to remove age-associated changes that could be driven by strong early life developmental processes. At Cayo Santiago some females exhibit signs of oestrus at that age (*81*), and males typically emigrate from their natal groups which confirms that they have entered late developmental phase (*82*).

Models testing the effect of age included sex, group membership, percent uniquely mapped reads, and a relatedness matrix as covariates. Models testing the effect of ELA included age as an additional covariate. To avoid confounding investigations of early life adversities with the current social environment, we also included current social status (methods described above) in ELA models. We binned cumulative ELA into individuals experiencing 0, 1, 2, or 3+ adversities in early life, as has been done in prior work (*8*). We tested the effect of each binary ELA variable on DNAm separately, as well as the effect of cumulative ELA. Models that failed to converge were excluded (mean ± SD = 2.7 ± 3.3%, range = 0.5-12.6% of tested sites across tissues for the effect of age; mean ± SD = 1.7 ± 0.9%, range = 0.6-4.5% of tested sites across tissues and ELAs for the effects of ELA).

To refine our model estimates, we used multivariate adaptive shrinkage implemented in *mashr* (*83*) which provides a statistical power gain by leveraging the sharing of information across conditions. Specifically, *mashr* uses the effect sizes and associated standard errors for the effect of a predictor at a given locus across conditions to 1) estimate pairwise condition-by-condition correlation and covariance across conditions, and 2) use the acquired information to refine the effect estimates and measures of significance (in the form of local false sign rate) using Bayes' theorem. Because *mashr* does not allow for missing values, we restricted further analysis to regions present in all tissues (157,609 regions from a maximum of 190,868 tested for age effects; mean ± SD = 157,402 ± 2,227, range = 152,840-158,948 regions tested for ELA effects), and excluded gonads which had half of the sample sizes of other tissues.

Data was processed as instructed on the mashr github tutorial https://stephenslab.github.io/mashr/articles/intro_mash.html, separately for age and ELA-effects and following the same procedure. Specifically, we defined a set of strongly *predictor*-associated regions (age: 15,069 regions achieving q-value < 0.01 in pqlseq and covered in all tissues; ELA: mean ± SD = 20,559.6 ± 1,340.8, range = 18,383-22,274 regions achieving a p-value < 0.01 in pqlseq and covered in all tissues). Then, we estimated background correlation by using estimate_null_correlation_simple() on the full data, and updated the full data and strong set using the estimated null covariance matrix. Data driven covariance was estimated on the strong set using cov_pca() and cov_flash(), and passed for extreme deconvolution to cov_ed(). Canonical covariance matrices were assessed on the full data using cov_canonical(). Finally, we calculated the posterior probabilities for the beta effects, and associated standard deviation and local false sign rates (LFSR) using mash() on the full data with data driven covariance structure passed to the Ulist= argument.

Regions were considered age- and ELA-associated for LFSR < 0.05 and < 0.1 respectively. We considered effects to be shared across tissues if their effect size was within a log-fold change of 2 to the tissue with the strongest effect. Therefore, regions significant in only one tissue and for which effect was not shared in another tissue were considered as uniquely age- or ELA-associated regions for that tissue. Regions significant in all 12 tissues were considered fully shared. All other regions were considered partially shared if they exhibited associated differences in between two and 11 tissues, or not associated if significant in no tissue. Pairwise sharing of effects were calculated using get_pairwise_sharing() from *mashr* with factor=0.5, lfsr_thresh = 0.05. The function calculates correlation of effect sizes between two tissues at all sites exhibiting significant changes in at least one of the tissues in the pair. We plotted the pairwise sharing using the package *corrplot* (85). For illustrative purposes, we also plotted percent methylation according to age and ELA for one tissue-specific and one fully-shared region (note that the significance was determined from binomial models accounting for differences in coverage).

<u>Enrichment for genomic features, chromatin marks and gene set annotations</u>
Genomic coordinates corresponding to gene bodies, promoters, and CpG islands were downloaded from UCSC Genome Browser (http://genome.ucsc.edu) for the rheMac10 genome (*59*). We downloaded chromatin marks chromHMM annotations available for 10 of our 12 tissues through ENCODE from the Roadmap Epigenomics project (downloaded from https://egg2.wustl.edu/roadmap/data/byFileType/chromhmmSegmentations/ChmmModels/core

Marks/jointModel/final/) and transposed to rhesus macaque genomic coordinates using liftOver. These included the E063 for adipose tissue, E066 for the liver, E080 for the adrenal, E086 for the kidney, E096 for the lung, E105 for the heart, E108 for the skeletal muscles, E112 for the thymus, E113 for the spleen, and E062 for PBMCs. All annotations were intersected with the set of all sites tested for age- and ELA-associated changes using the bedtools intersect function (*84*). We tested for enrichment of chromatin marks within our significantly associated sites in each tissue using a Fisher's exact test with Benjamini-Hochberg correction for multiple hypothesis testing (*85*).

Lastly, we aimed to test whether tissues with particular cell types or biological functions exhibited stronger ELA effects. To do so, we categorized tissues into those possessing long (> 180 days) versus short (< 180 days) cell lifespans, as this has proven to be an important determinant of cell-type specific aging in the mouse (*18*). We used cell lifespan data for the most abundant cell type within each tissue determined from human cells for all tissues except for the spleen, for which cell lifespan data was extrapolated from the mouse (*86*). We fit a single linear model, including cell lifespan (short/long) and biological system (immune/endocrine/metabolic/other) as predictor variables, ELA variable as a covariate, and the absolute value of the ELA effect size as the response variable.

Prediction of tissue-specific DNAm age with elastic net regression

We built tissue-specific elastic net regression models on chronological age using *glmnet* (*58*). Regions mapping to the X chromosome were removed to avoid strong sex bias in model prediction, and restricted to regions covered in all tissues because *glmnet* does not allow missing value. We applied a classic log transformation on age before maturity (set to 5 years old) because methylation profiles tend to change faster in development (*87*). We used a leave-one-out validation to achieve unbiased prediction for each sample. The model was computed on the training set using cv.glmnet and the penalty parameter lambda was optimized via internal 10-fold cross-validation. The prediction for the test sample was obtained by applying the model with the function predict(). Alpha was chosen a posteriori to minimize the average MSE in a given tissue. Model performance was assessed by calculating the Pearson's correlation coefficient and Median Average Error between chronological and predicted age. Differences in clocks' performance were not explained by technical artefacts such as sample size, age range, or variance of individuals' age in the respective tissue set (fig. S9).

Final clock coefficients were extracted from models fitted on all samples in a given tissue with alpha set to the value defined via the leave-one-out procedure. To better understand which regions were more susceptible to be selected by the clock, we assessed the correlation between percent methylation and age in regions selected and those omitted from the clocks (fig. S8).

Tissue-specific age deviations

Predicted age in mature individuals (i.e., >3 years old) was regressed on chronological age with the function lm(), and residuals were extracted with the function residuals() to express an individual deviation from the expected DNAm. Linear regression fits were assessed and validated by inspecting the distribution of residuals with a QQplot. To exclude that DNAm deviations could be associated with technical or cohort effects, we tested for an association between DNAm and several covariates, including sample sizes, age range, and age deviation for individuals available for each tissue set (figs. S9 and S10). We tested for greater consistency in individual age deviations based on the assumption that it would be smaller than at random by permuting tissue-specific deviations across individuals, and calculating average within-individual variance on the randomized set, done 10,000. This procedure generated a distribution of random within-individual heterogeneity matched to our empirical data and compared whether the observed average within-individual variance fell within the randomized distribution. As permutations can fail to fully account for confounding factors (*88*)—for example sex or age—, we generated an unbiased estimation of the proportion of variance explained by individual ID by fitting a linear mixed model on age predictions including sex, tissue type, group, and chronological age as fix effects terms, and individual ID as a random effects term https://bbolker.github.io/mixedmodels-misc/glmmFAQ.html#testing-significance-of-random-effects. A p-value for the comparison of models with and without the random effects term for individual ID was calculated using exactLRT() from the package *RLRsim* (*89*).

Next, we aimed to investigate correlations among tissue-specific age-deviations to identify potential multi-tissue clusters of age-associated changes. As missing values were not allowed, we imputed missing samples in 190 individuals using the function mice(m = 5, method = "pmm", maxit = 35) from the package *mice* (*90*), and extracted the first imputed run after assessing imputation accuracy and convergence. Individuals were dropped if more than ⅓ of their tissue values were imputed, leaving 179 individuals in the final set. Pairwise comparisons of age deviations were calculated using cor() and plotted with corrplot(order = 'hclust', hclust.method = "complete"). We performed hierarchical clustering on the distance matrix using dis(method=euclidean) and hclust(method=complete) and plotted with ComplexHeatmap (*91*). To test whether individuals would fall into different clusters i.e., ageotypes, we determined the most parsimonious number of clusters with *useful* FitKMeans() and PlotHartigan(), and applied kmeans clustering with *factoextra* eclust() (fig. S11 and Supplemental results).

<u>Contribution of early life adversity to age-associated differences in DNA methylation</u>

We investigated how ELA contributes to aging-related differences in DNAm in three distinct ways. First, we determined whether ELA predicts an individual's deviation from their chronological age as determined via our tissue-specific epigenetic clocks. We fit linear mixed-effect models testing for 1) the additive effect of each ELA on residual age (i.e., age deviation) controlling for tissue, and 2) the nested effect of ELA within each tissue. All models included sex and chronological age as covariates and individual ID as a random effect to account for repeated measures. We compared additive and nested models using likelihood ratio tests, and report results from the additive models, which were consistently supported by lower AIC values (dAIC > 2). To further investigate tissue-specific associations between ELA and age deviation, we fit linear models testing for the additive effect of each ELA within each tissue separately, controlling for sex and chronological age. Second, to understand whether ELA targets similar regions as those associated with age, we tested whether ELA and age-associated sites overlap more than would be expected by chance using Fisher's exact test. Third, to understand whether ELA effects may exacerbate DNAm changes that occur with age, we limited our analysis to sites that were significantly associated with both age and ELA and binned them into those exhibiting a positive effect (increased methylation with increased age or ELA exposure) and those exhibiting a negative effect (decreased methylation with increased age or ELA exposure). We then tested whether age- and ELA-associated sites were more likely to have the same direction or effect than would be expected by chance using Fisher's exact test. For all analyses, we used the Benjamini-Hochberg correction for multiple hypothesis testing (*85*).

### Supplemental Results

The aging methylome landscape across tissues

Detailed assessment of the proportion of sites increasing versus decreasing in methylation with age revealed important differences across tissues. We found that the thymus, pituitary, spleen, and whole blood exhibited a strong hypomethylation bias with age (ratio for the number of hypermethylated/hypomethylated sites ranging from 0.07 to 0.23). The skeletal muscle, and adrenal showed moderate hypomethylation biases (ratios 0.79 and 0.80). While the liver and omental showed a slight hypermethylation bias with age (ratios 1.01 and 1.17), the latter was much more pronounced in the kidney, thyroid, heart, and lung (ratio 1.59 to 2.97).

Enrichment tests showed that tissue-specific markers are more likely than expected by chance to exhibit age-associated changes. However, this enrichment was not significantly different from results obtained following the permutations of tissue-specific markers in the adrenal, skeletal muscle, and pituitary. This would suggest that markers and age-associated sites broadly overlap but that tissue-specific markers are not more likely to exhibit age-associated changes in the associated tissue, which is consistent with previous reports of limited overlap between sites involved in tissue differentiation and aging (*40*, *92*).

The aging methylome of the thymus

The thymus exhibited a marked hypomethylation bias with age, consistent with previous studies in macaques (*29*, *93*) and in T-cells with age in humans (*94*). Notably, thymus profiles were more similar to the adipose tissue than to other samples, and this similarity appeared to increase with age (fig. S2), which is consistent with adipose infiltration occurring during involution. Moreover, we note several concordant observations with previous research, for example the high methylation level of the *ngf* gene (average percent methylation = 0.92), a transcription factor central to epithelial cell proliferation and to the RAP1 signaling pathway, and which methylation levels are decreased by anti-aging stem cell therapy (*93*). Age-associated differences at regions overlapping several genes in our dataset were also previously reported (e.g., hypomethylation of *gad2*, *fgf2*, *fcn3*, *adamdec1*, *f13a1*, *c3* (*29*)), although we did not observe an hypermethylation of *foxn1*, which has been suggested as a marker of thymic involution (*95*) but had high average methylation levels in our data (average percent methylation = 0.74 across six regions).

Sharing of age-associated effects across 10 tissues

Leaving the thymus and pituitary aside, an age-DMR was significant at LFSR < 0.05 in 2.3 ± 2.2 tissues on average (median = 1), with "tissue-shared" effects in 3.0 ± 2.5 tissues (median = 2) (fig. S6). Therefore, the number of consistent DNAm changes across tissues slightly increased when omitting the thymus and pituitary, although most changes remained largely independent across tissues.

Age predictions in external dataset for whole blood

To test the reliability of the DNA methylation clock generated using the cross-sectional data from Cayo rhesus macaques, we applied the whole blood to a new dataset of samples collected longitudinally on Cayo Santiago, and that were not included in the present study (n = 642). We used the alpha value minimizing MSE in the cross-sectional sample. Age predictions exhibited a Pearson's correlation = 0.76, with a MAE = 2.18 (table S11). Although lower than on the original cross-sectional data, this performance remains high showing that the clock generated captures well methylome information related to aging in this population.

<u>DNA methylation profiles across tissues quantifies within-individual heterogeneity of aging</u>
Using tissue-specific age deviations (fig. 3F), we attempted to identify patterns of covariance across individuals (also named "ageotypes") using kmeans clustering. However, fviz_silhouette() revealed that several individuals could not be unambiguously classified (fig. S11A). Furthermore, the clustering procedure separated individuals into two groups, with one group exhibiting positive age-devations across tissues (i.e., older than expected biological ages) whereas the other group gathered individuals with negative age-deviations across tissues (i.e., younger than expected biological ages) (fig. S11, B and C). Thus, we could not identify ageotypes in this study sample, although we acknowledge that our sample size is smaller in comparison to previous studies which suggested the existence of ageotypes in humans (*96*).

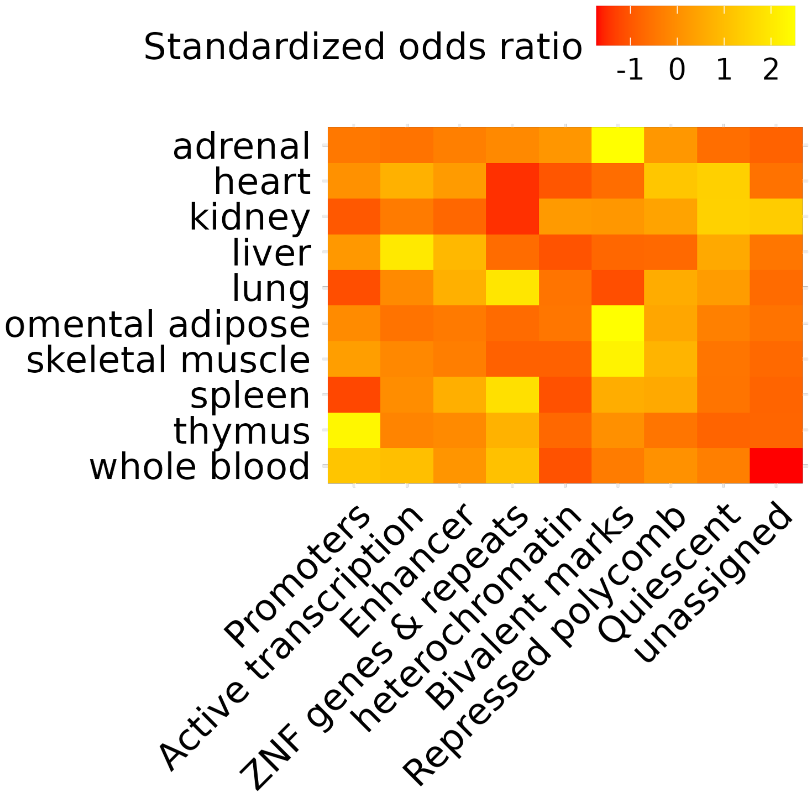


**Fig. S1. Chromatin annotation of organ-specific hypermethylated markers.** Odds ratios were standardized by z-transformation across annotations within organs for visualization.

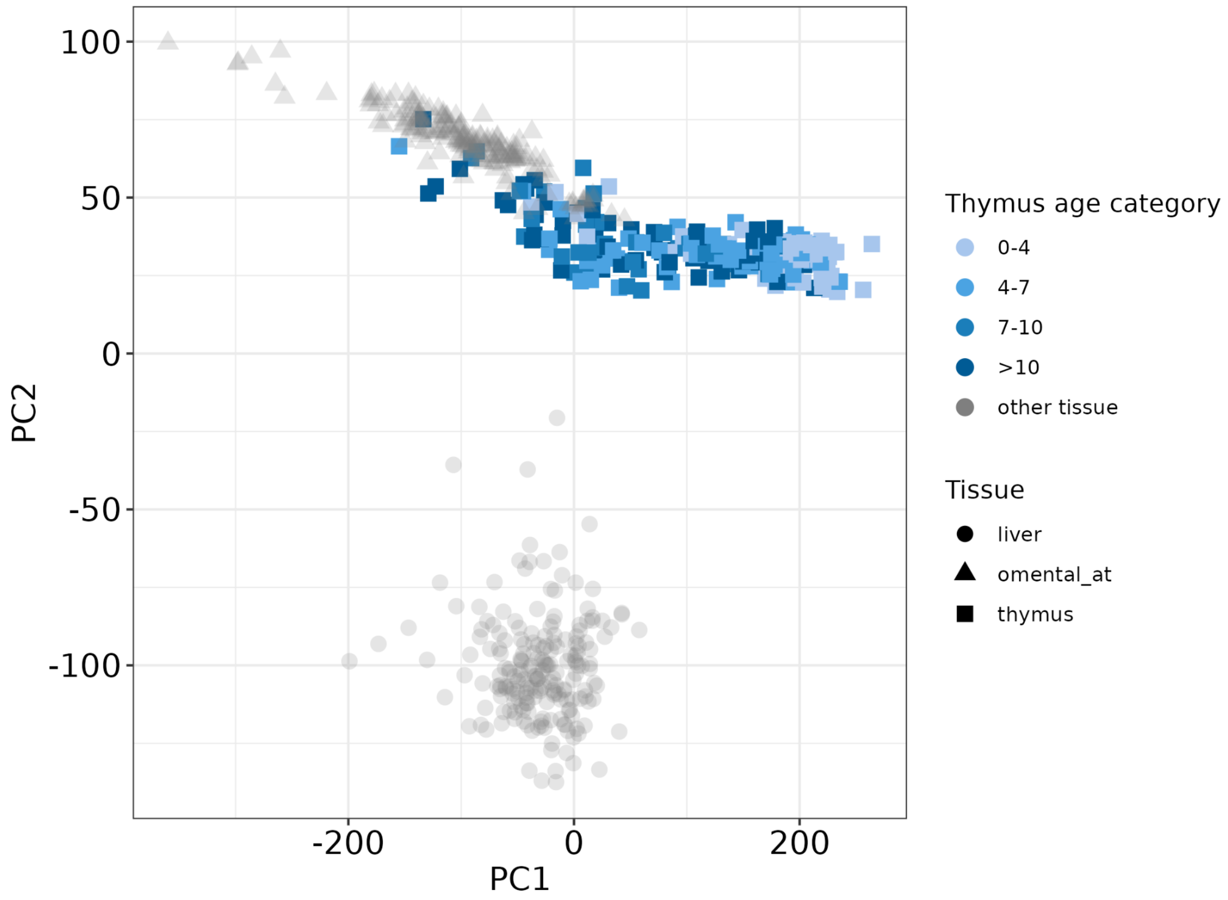


**Fig. S2. Thymus exhibit age-associated transition towards adipose-like methylation profiles.** The PCA was performed using 47,699 regions with complete coverage across 610 samples (N thymus = 224, omental adipose = 185, liver = 201). Thymus samples exhibit age-associated loadings on PC1 (explaining 26% of variance) with a transition towards omental adipose samples, compatible with the known adipose infiltration occurring during thymus involution. The liver was chosen as an "outgroup" tissue.

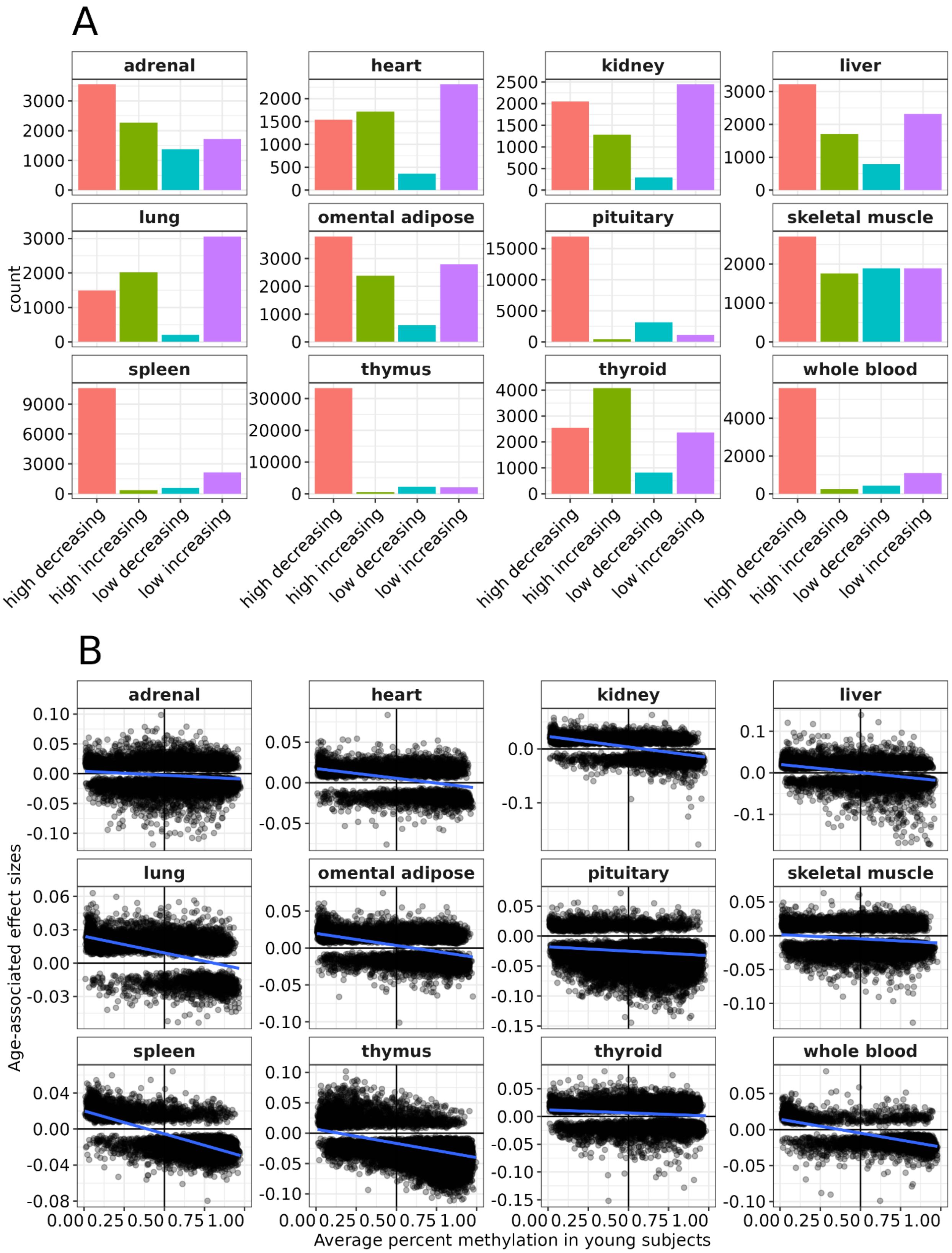
A
adrenal
heart
kidney
liver
lung
omental adipose
pituitary
skeletal muscle
spleen
thymus
thyroid
whole blood
count
high decreasing
high increasing
low decreasing
low increasing
B
adrenal
heart
kidney
liver
lung
omental adipose
pituitary
skeletal muscle
spleen
thymus
thyroid
whole blood
Age-associated effect sizes
0.00 0.25 0.50 0.75 1.00
Average percent methylation in young subjects

**Fig. S3. CpG methylation levels are closer to 0.5 with increasing age.** (**A**) Average percent methylation in young individuals was categorized as higher or lower than 0.5. Only sites with LFSR < 0.05 for the effect of age were included for both analyses. (**B**) Negative correlations between average percent methylation in young individuals (< 6 years of age) and age-associated effect sizes. Fitted lines were plotted from linear regressions with geom_smooth(method = "lm") in *ggplot2*.

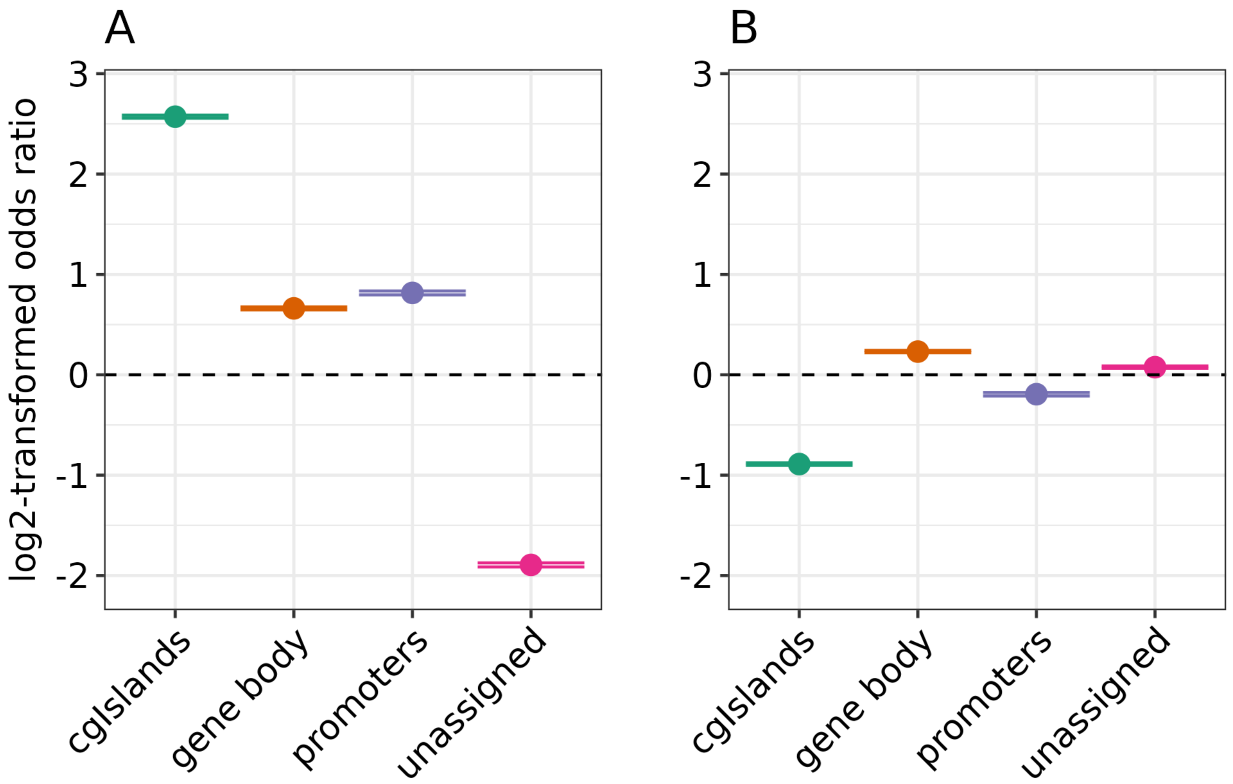


**Fig. S4. Enrichment tests for functional annotations.** Sites were divided as (**A**) hypermethylated and (**B**) hypomethylated with age.

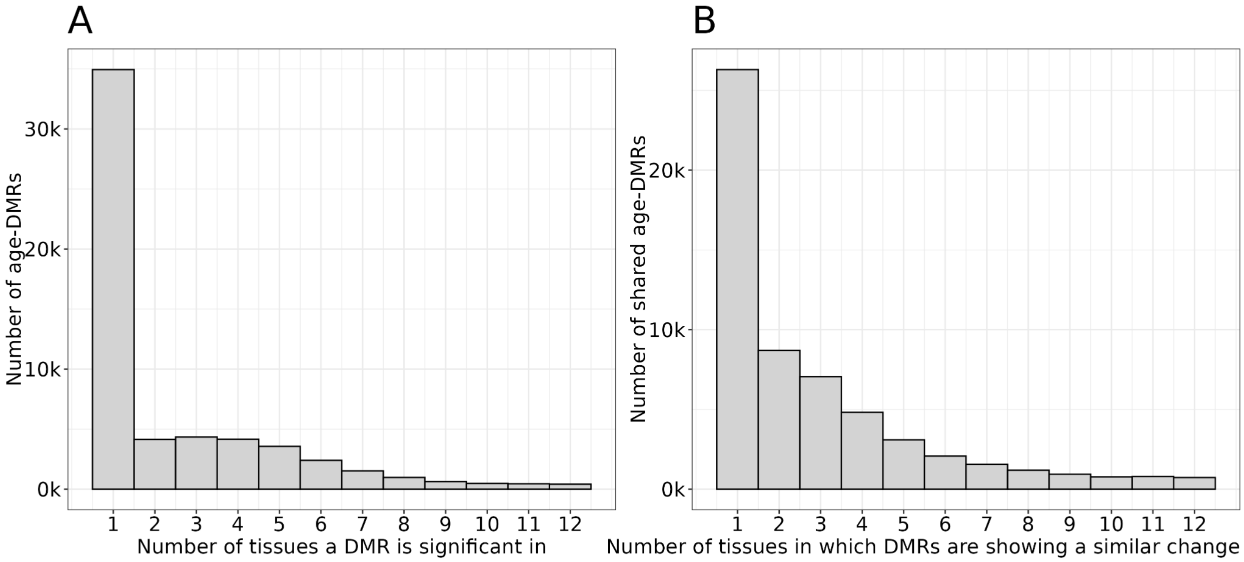


**Fig. S5. Cross-tissue sharing of methylation age differences.** Number of tissues in which DMRs (**A**) exhibited age-associated changes at a LFSR < 0.05, and (**B**) shared effect sizes of similar magnitude.

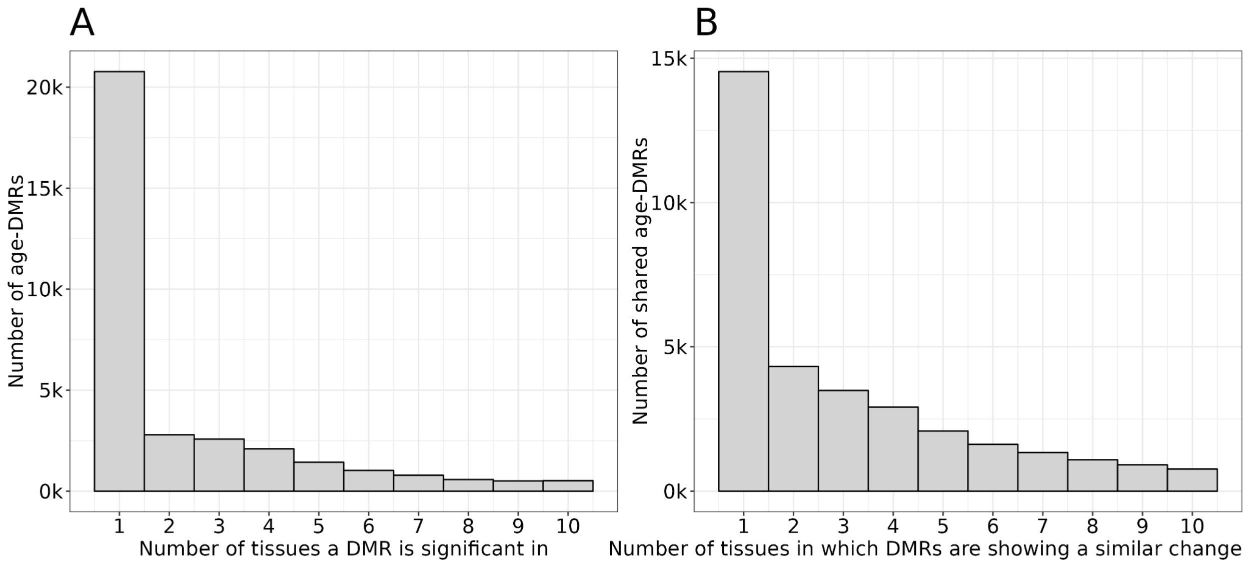


**Fig. S6. Robustness test of cross-tissue sharing of methylation age differences.** Excluding the thymus and pituitary showing extensive age-associated DNA methylation only marginally increased the number of tissues in which DMRs (**A**) exhibited age-associated changes at a LFSR < 0.05, and (**B**) shared effect sizes of similar magnitude.

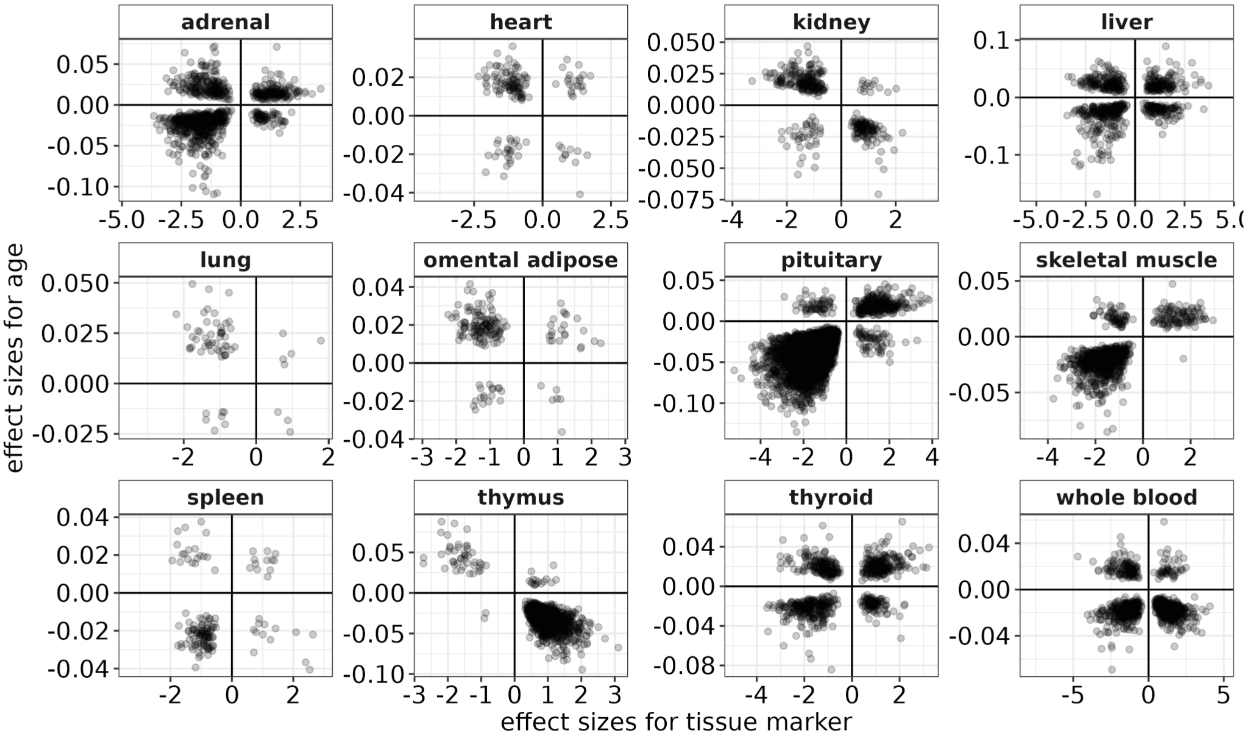


**Fig. S7. Tissue-specific markers exhibiting age-associated differences in percent methylation.** On the x-axis, the effect sizes are negative for hypomethylated markers and positive for hypermethylated markers. Corresponding effect sizes for increasing and decreasing levels of methylation with age are shown on the y-axis. Sites falling into the upper-left and lower-right quadrants are losing specificity with age, whereas those falling into the lower-right and upper-left quadrants are gaining specificity with age.

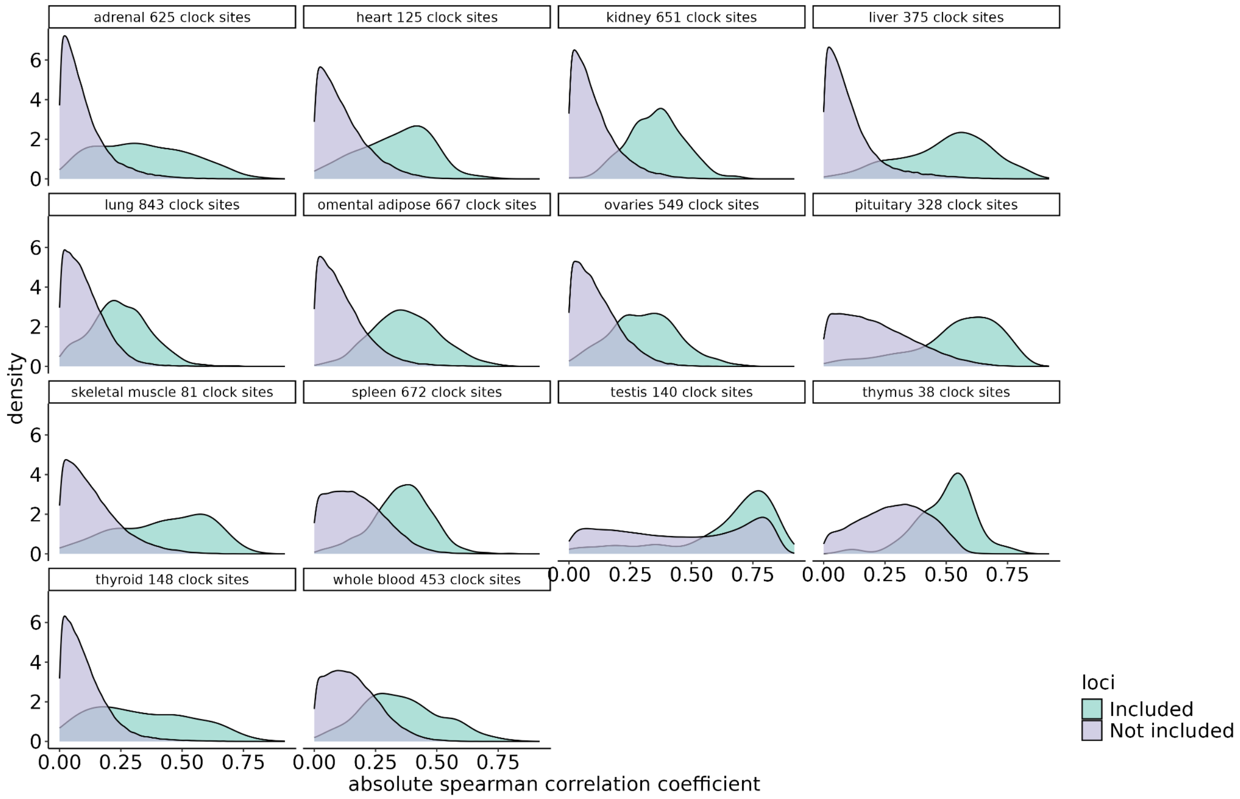


**Fig. S8. Correlation of methylation levels of clocks' regions and age.** Correlation of methylation levels with age are shown for regions included or not in the clock. Percent methylation correlated with age significantly more at regions included in the clock than at regions not included in whole blood, omental adipose, and pituitary (Fisher's z-transformation of the correlation coefficients, p-value < 0.05), but not in other tissues.

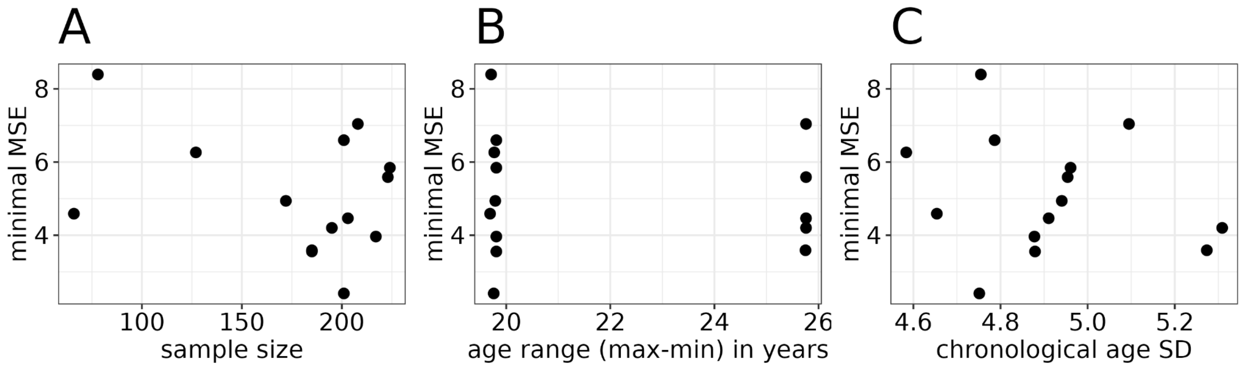


**Fig. S9. Tissue-specific clock performance.** Median Squared Error (MSE) is unrelated to (**A**) sample size, (**B**) age range, or (**C**) variability in ages across individuals.

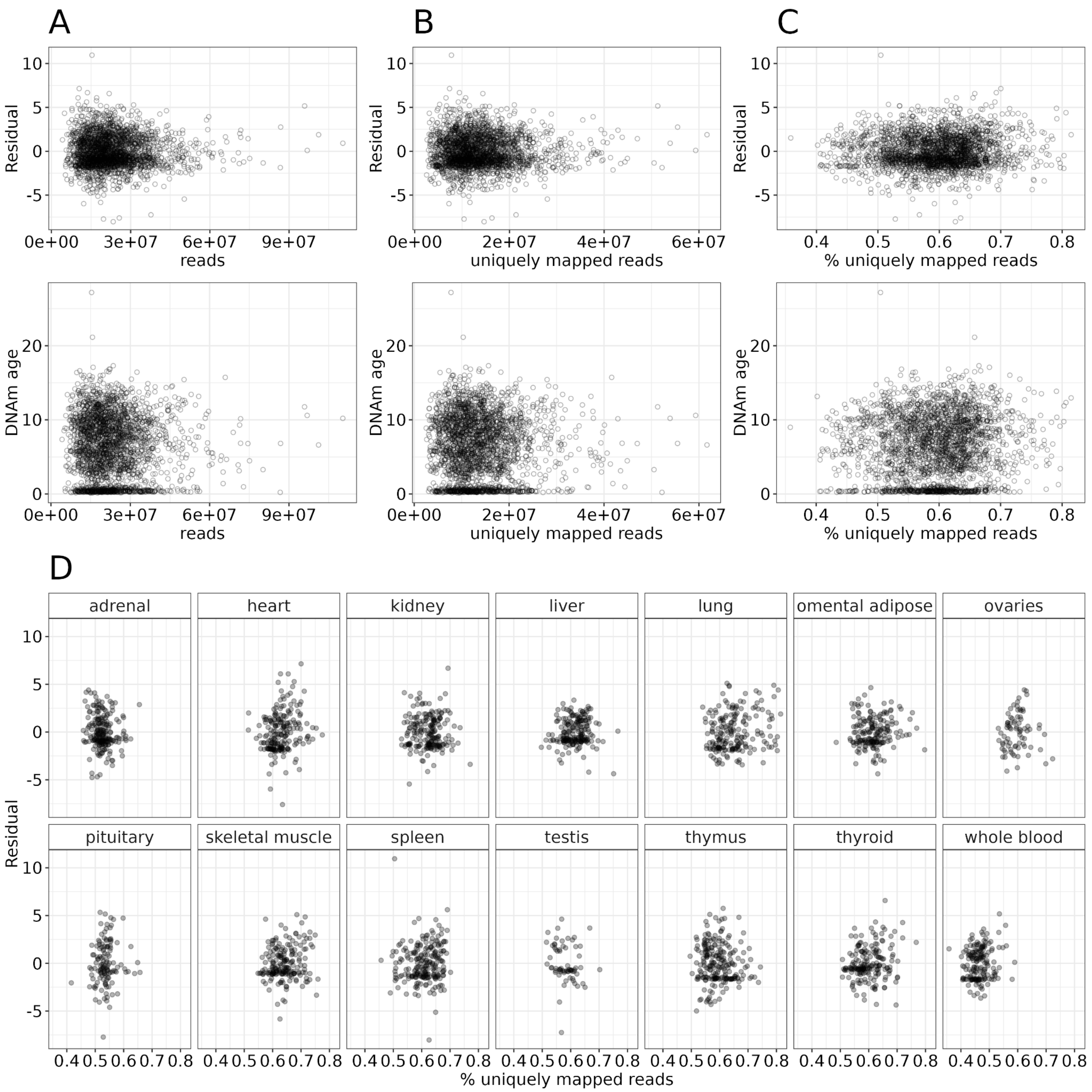


**Fig. S10. Technical correlates of DNAm age prediction**. (**A**) DNAm age predictions are unbiased with respect to sequencing depth (i.e., total reads), (**B**) number of uniquely mapped reads, or (**C**) the percentage of uniquely mapped reads to total reads. (**D**) Breakdown of residual against percent uniquely mapped reads per tissue.

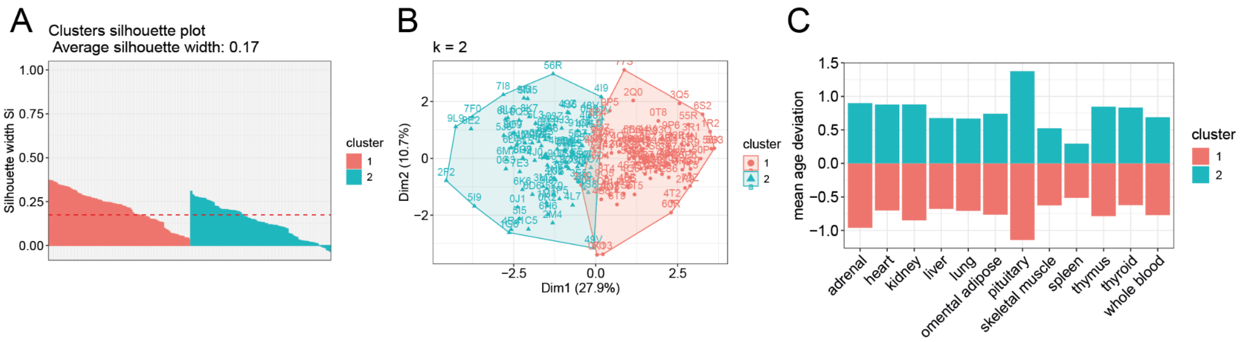


**Fig. S11. Kmeans clustering of age deviations.** (**A**) The silhouette diagnostic plot shows that some individuals could not be assigned unambiguously (negative scores on the y-axis). (**B**) Unsupervised clustering assigned individuals across two groups. (**C**) Clusters exhibit on average consistent age-acceleration or deceleration across tissues.

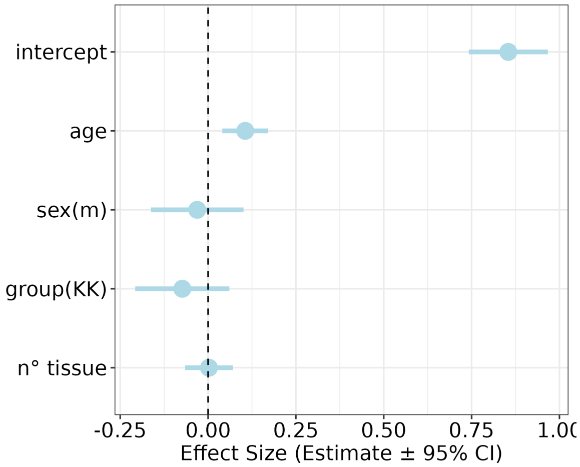


**Fig. S12. Predictors of within-individual heterogeneity.** Within-individual heterogeneity increases with age but does not differ between females and males. Within-individual heterogeneity is measured by calculating the variance in DNAm age across tissues for each individual. The number of tissues available was included as a control.

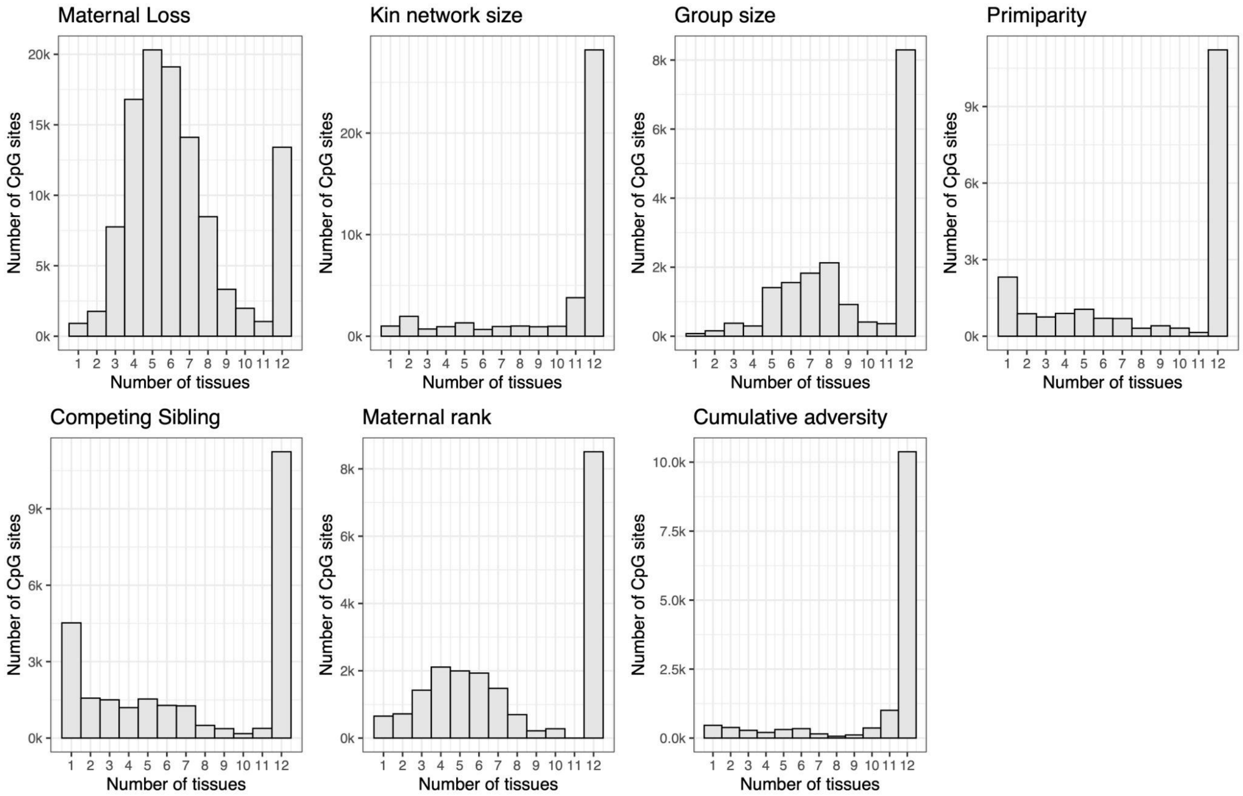


**Fig. S13. The intersection of CpG sites associated with each adversity across tissues following the effect size sharing analysis.** Epigenetic response to any given ELA is generally coordinated across multiple tissues.

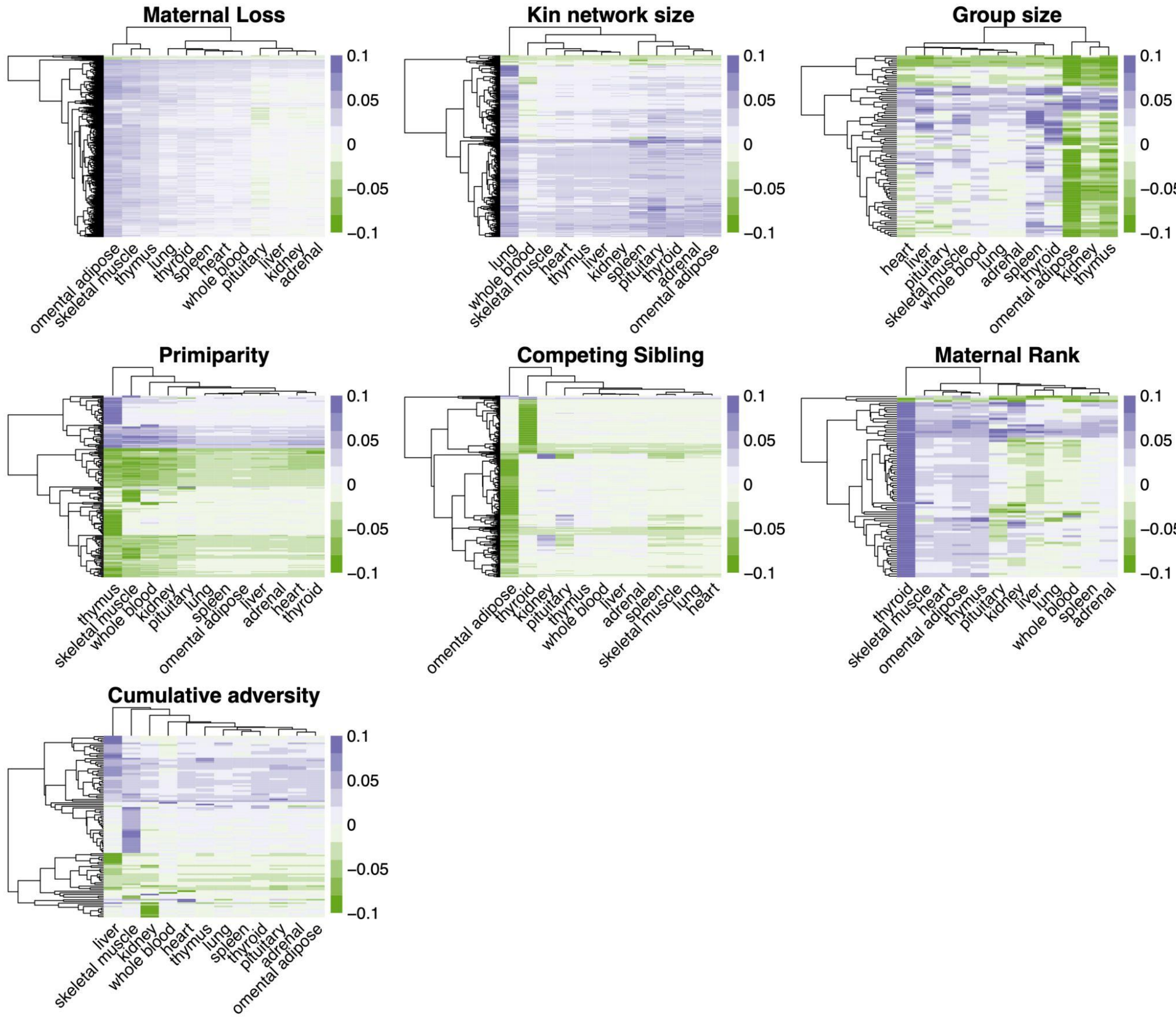


**Fig. S14. Hierarchical clustering of sites associated with each ELA**. Cells are colored by effect size (positive = increased methylation, negative = decreased methylation), limited to sites that did not have shared effects across all 12 tissues.

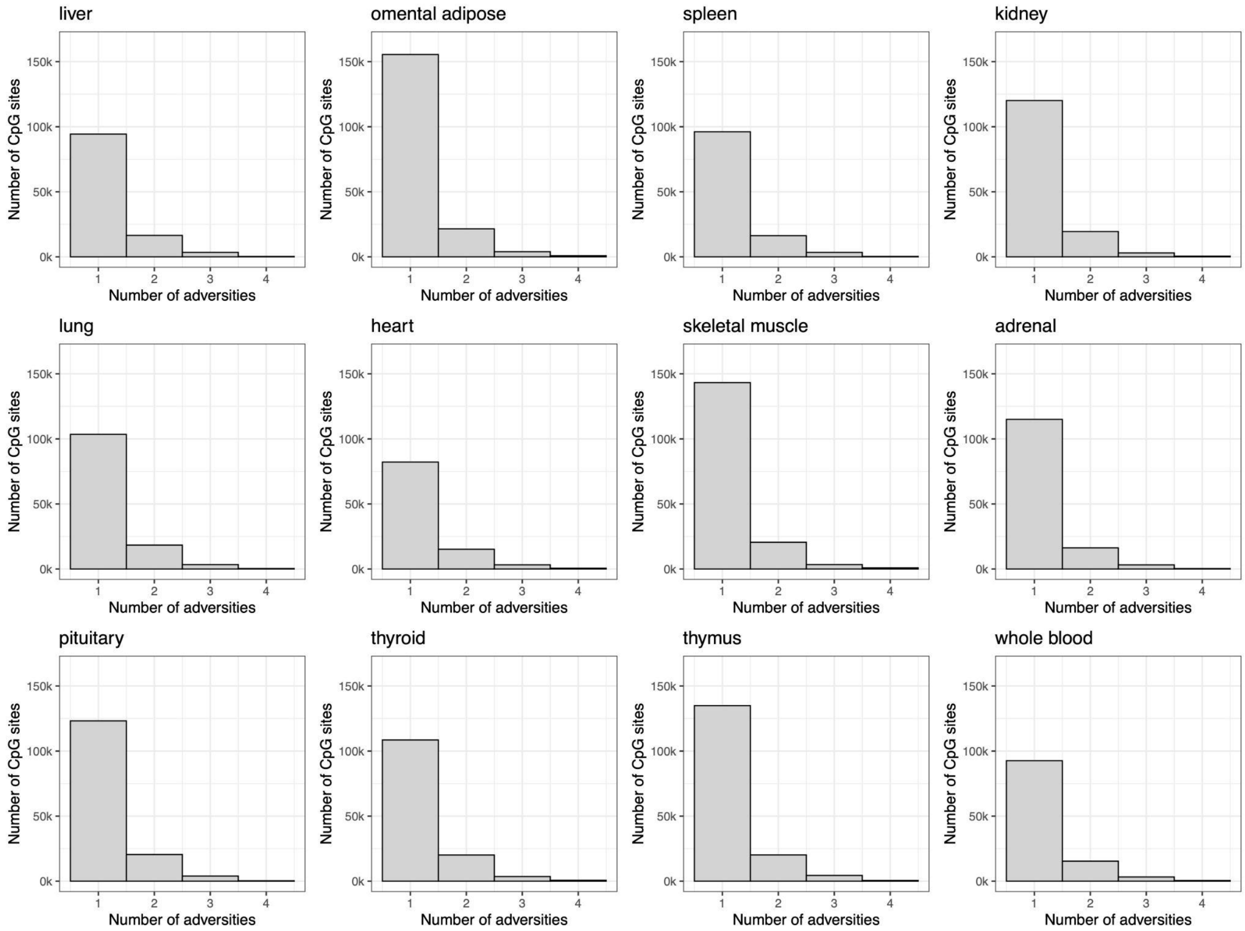


**Fig. S15. The intersection of CpG sites associated with each ELA across tissues.**

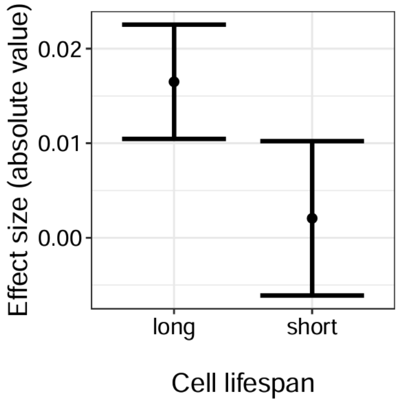


**Fig. S16. Stronger ELA effects in tissues with long versus short-lived cell types.** Linear model results testing for the effect of cell lifespan (short versus long) in predicting the strength of ELA effect. Points represent model predictions and error bars represent the 0.95 confidence interval.

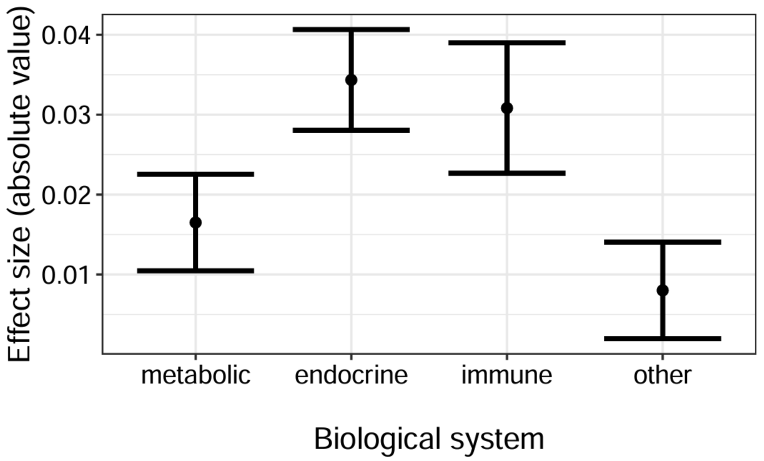


**Fig. S17. Stronger ELA effects in endocrine and immune tissues.** Linear model results testing for the effect of biological systems in predicting the strength of ELA effect. Points represent model predictions and error bars represent the 0.95 confidence interval.

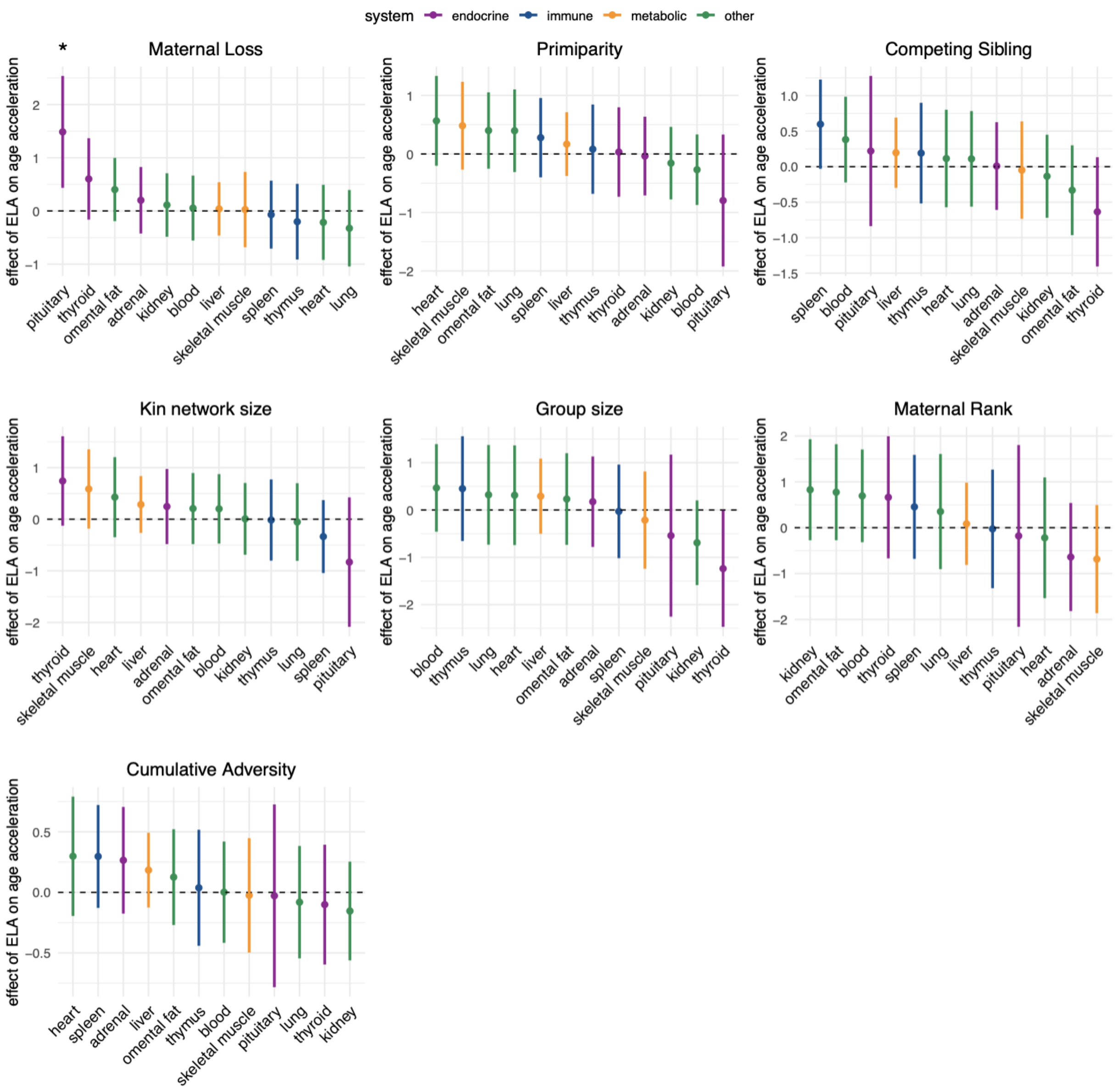


**Fig. S18. Tissue-dependent effects of ELAs on age deviation.** Linear mixed-effect model results testing for the effect of each adversity on age deviation (biological - chronological age) as determined by our tissue-specific epigenetic clocks. Points represent the model prediction, bars represent the 0.95 confidence interval, and colors represent which biological system each tissue is attributed to.

17. B. E. Stranger, L. E. Brigham, R. Hasz, M. Hunter, C. Johns, M. Johnson, G. Kopen, W. F. Leinweber, J. T. Lonsdale, A. McDonald, B. Mestichelli, K. Myer, B. Roe, M. Salvatore, S. Shad, J. A. Thomas, G. Walters, M. Washington, J. Wheeler, J. Bridge, B. A. Foster, B. M. Gillard, E. Karasik, R. Kumar, M. Miklos, M. T. Moser, S. D. Jewell, R. G. Montroy, D. C. Rohrer, D. R. Valley, D. A. Davis, D. C. Mash, S. E. Gould, P. Guan, S. Koester, A. R. Little, C. Martin, H. M. Moore, A. Rao, J. P. Struewing, S. Volpi, K. D. Hansen, P. F. Hickey, L. F. Rizzardi, L. Hou, Y. Liu, B. Molinie, Y. Park, N. Rinaldi, L. Wang, N. Van Wittenberghe, M. Claussnitzer, E. T. Gelfand, Q. Li, S. Linder, R. Zhang, K. S. Smith, E. K. Tsang, L. S. Chen, K. Demanelis, J. A. Doherty, F. Jasmine, M. G. Kibriya, L. Jiang, S. Lin, M. Wang, R. Jian, X. Li, J. Chan, D. Bates, M. Diegel, J. Halow, E. Haugen, A. Johnson, R. Kaul, K. Lee, M. T. Maurano, J. Nelson, F. J. Neri, R. Sandstrom, M. S.

Fernando, C. Linke, M. Oliva, A. Skol, F. Wu, J. M. Akey, A. P. Feinberg, J. B. Li, B. L. Pierce, J. A. Stamatoyannopoulos, H. Tang, K. G. Ardlie, M. Kellis, M. P. Snyder, S. B. Montgomery, eGTEx Project, Enhancing GTEx by bridging the gaps between genotype, gene expression, and disease. *Nat. Genet.* **49**, 1664–1670 (2017).

18. M. J. Zhang, A. O. Pisco, S. Darmanis, J. Zou, Mouse aging cell atlas analysis reveals global and cell type-specific aging signatures. *eLife* **10**, e62293 (2021).

19. M. J. Kessler, R. G. Rawlins, A 75-Year Pictorial History of the Cayo Santiago Rhesus Monkey Colony. *Am. J. Primatol.* **78**, 6–43 (2016).

20. C. López-Otín, M. A. Blasco, L. Partridge, M. Serrano, G. Kroemer, Hallmarks of aging: An expanding universe. *Cell* **186**, 243–278 (2023).

21. M. V. C. Greenberg, D. Bourc'his, The diverse roles of DNA methylation in mammalian development and disease. *Nat. Rev. Mol. Cell Biol.* **20**, 590–607 (2019).

22. A. Haghani, C. Z. Li, T. R. Robeck, J. Zhang, A. T. Lu, J. Ablaeva, V. A. Acosta-Rodríguez, D. M. Adams, A. N. Alagaili, J. Almunia, A. Aloysius, N. M. S. Amor, R. Ardehali, A. Arneson, C. S. Baker, G. Banks, K. Belov, N. C. Bennett, P. Black, D. T. Blumstein, E. K. Bors, C. E. Breeze, R. T. Brooke, J. L. Brown, G. Carter, A. Caulton, J. M. Cavin, L. Chakrabarti, I. Chatzistamou, A. S. Chavez, H. Chen, K. Cheng, P. Chiavellini, O.-W. Choi, S. Clarke, J. A. Cook, L. N. Cooper, M.-L. Cossette, J. Day, J. DeYoung, S. Dirocco, C. Dold, J. L. Dunnum, E. E. Ehmke, C. K. Emmons, S. Emmrich, E. Erbay, C. Erlacher-Reid, C. G. Faulkes, Z. Fei, S. H. Ferguson, C. J. Finno, J. E. Flower, J.-M. Gaillard, E. Garde, L. Gerber, V. N. Gladyshev, R. G. Goya, M. J. Grant, C. B. Green, M. B. Hanson, D. W. Hart, M. Haulena, K. Herrick, A. N. Hogan, C. J. Hogg, T. A. Hore, T. Huang, J. C. Izpisua Belmonte, A. J. Jasinska, G. Jones, E. Jourdain, O. Kashpur, H. Katcher, E. Katsumata, V. Kaza, H. Kiaris, M. S. Kobor, P. Kordowitzki, W. R. Koski, M. Krützen, S. B. Kwon, B. Larison, S.-G. Lee, M. Lehmann, J.-F. Lemaître, A. J. Levine, X. Li, C. Li, A. R. Lim, D. T. S. Lin, D. M. Lindemann, S. W. Liphardt, T. J. Little, N. Macoretta, D. Maddox, C. O. Matkin, J. A. Mattison, M. McClure, J. Mergl, J. J. Meudt, G. A. Montano, K. Mozhui, J. Munshi-South, W. J. Murphy, A. Naderi, M. Nagy, P. Narayan, P. W. Nathanielsz, N. B. Nguyen, C. Niehrs, B. Nyamsuren, J. K. O'Brien, P. O. Ginn, D. T. Odom, A. G. Ophir, S. Osborn, E. A. Ostrander, K. M. Parsons, K. C. Paul, A. B. Pedersen, M. Pellegrini, K. J. Peters, J. L. Petersen, D. W. Pietersen, G. M. Pinho, J. Plassais, J. R. Poganik, N. A. Prado, P. Reddy, B. Rey, B. R. Ritz, J. Robbins, M. Rodriguez, J. Russell, E. Rydkina, L. L. Sailer, A. B. Salmon, A. Sanghavi, K. M. Schachtschneider, D. Schmitt, T. Schmitt, L. Schomacher, L. B. Schook, K. E. Sears, A. W. Seifert, A. B. A. Shafer, A. V. Shindyapina, M. Simmons, K. Singh, I. Sinha, J. Slone, R. G. Snell, E. Soltanmohammadi, M. L. Spangler, M. Spriggs, L. Staggs, N. Stedman, K. J. Steinman, D. T. Stewart, V. J. Sugrue, B. Szladovits, J. S. Takahashi, M. Takasugi, E. C. Teeling, M. J. Thompson, B. Van Bonn, S. C. Vernes, D. Villar, H. V. Vinters, H. Vu, M. C. Wallingford, N. Wang, G. S. Wilkinson, R. W. Williams, Q. Yan, M. Yao, B. G. Young, B. Zhang, Z. Zhang, Y. Zhao, P. Zhao, W. Zhou, J. A. Zoller, J. Ernst, A. Seluanov, V. Gorbunova, X. W. Yang, K. Raj, S. Horvath, DNA methylation networks underlying mammalian traits. *Science* **381**, eabq5693 (2023).

30. N. Loyfer, J. Magenheim, A. Peretz, G. Cann, J. Bredno, A. Klochendler, I. Fox-Fisher, S. Shabi-Porat, M. Hecht, T. Pelet, J. Moss, Z. Drawshy, H. Amini, P. Moradi, S. Nagaraju, D. Bauman, D. Shveiky, S. Porat, U. Dior, G. Rivkin, O. Or, N. Hirshoren, E. Carmon, A. Pikarsky, A. Khalaileh, G. Zamir, R. Grinbaum, M. Abu Gazala, I. Mizrahi, N. Shussman, A. Korach, O. Wald, U. Izhar, E. Erez, V. Yutkin, Y. Samet, D. Rotnemer Golinkin, K. L. Spalding, H. Druid, P. Arner, A. M. J. Shapiro, M. Grompe, A. Aravanis, O. Venn, A. Jamshidi, R. Shemer, Y. Dor, B. Glaser, T. Kaplan, A DNA methylation atlas of normal human cell types. *Nature* **613**, 355–364 (2023).

46. N. Mariani, A. Borsini, C. A. M. Cecil, J. F. Felix, S. Sebert, A. Cattaneo, E. Walton, Y. Milaneschi, G. Cochrane, C. Amid, J. Rajan, J. Giacobbe, Y. Sanz, A. Agustí, T. Sorg, Y. Herault, J. Miettunen, P. Parmar, N. Cattane, V. Jaddoe, J. Lötjönen, C. Buisan, M. A. G. Ballester, G. Piella, J. L. Gelpi, F. Lamers, B. W. J. H. Penninx, H. Tiemeier, M. von Tottleben, R. Thiel, K. F. Heil, M.-R. Järvelin, C. Pariante, I. M. Mansuy, K. Lekadir, Identifying causative mechanisms linking early-life stress to psycho-cardio-metabolic multi-morbidity: The EarlyCause project. *PLOS ONE* **16**, e0245475 (2021).

53. N. N. Snyder-Mackler, Reduced Representation Bisulfite Sequencing (RRBS) with NEB Reagents. (2023).

59. W. C. Warren, R. A. Harris, M. Haukness, I. T. Fiddes, S. C. Murali, J. Fernandes, P. C. Dishuck, J. M. Storer, M. Raveendran, L. W. Hillier, D. Porubsky, Y. Mao, D. Gordon, M. R. Vollger, A. P. Lewis, K. M. Munson, E. DeVogelaere, J. Armstrong, M. Diekhans, J. A. Walker, C. Tomlinson, T. A. Graves-Lindsay, M. Kremitzki, S. R. Salama, P. A. Audano, M. Escalona, N. W. Maurer, F. Antonacci, L. Mercuri, F. A. M. Maggiolini, C. R. Catacchio, J. G. Underwood, D. H. O'Connor, A. D. Sanders, J. O. Korbel, B. Ferguson, H. M. Kubisch, L. Picker, N. H. Kalin, D. Rosene, J. Levine, D. H. Abbott, S. B. Gray, M. M. Sanchez, Z. A. Kovacs-Balint, J. W. Kemnitz, S. M. Thomasy, J. A. Roberts, E. L. Kinnally, J. P. Capitanio, J. H. P. Skene, M. Platt, S. A. Cole, R. E. Green, M. Ventura, R. W. Wiseman, B. Paten, M. A. Batzer, J. Rogers, E. E. Eichler, Sequence diversity analyses of an improved rhesus macaque genome enhance its biomedical utility. *Science* **370**, eabc6617 (2020).

96. R. Sehgal, M. Meer, A. H. Shadyab, R. Casanova, J. E. Manson, P. Bhatti, E. M. Crimmins, T. L. Assimes, E. A. Whitsel, A. T. Higgins-Chen, M. Levine, Systems Age: A single blood

methylation test to quantify aging heterogeneity across 11 physiological systems. bioRxiv [Preprint] (2023). https://doi.org/10.1101/2023.07.13.548904.
